# Identification of SLC45A4 as a pain gene encoding a neuronal polyamine transporter

**DOI:** 10.1101/2024.09.25.615046

**Authors:** Steven J Middleton, Sigurbjörn Markússon, Mikael Åkerlund, Justin C. Deme, Mandy Tseng, Wenqianglong Li, Sana R Zuberi, Gabriel Kuteyi, Peter Sarkies, Georgios Baskozos, Jimena Perez-Sanchez, Harry L Hébert, Sylvanus Toikumo, Adham Farah, Susan Maxwell, Yin Y Dong, Henry R Kranzler, John E Linley, Blair H Smith, Susan M. Lea, Joanne L Parker, Valeriya Lyssenko, Simon Newstead, David L Bennett

## Abstract

Polyamines are regulatory metabolites with key roles in transcription, translation, cell signalling and autophagy^1^. They are implicated in multiple neurological disorders including stroke, epilepsy and neurodegeneration and can regulate neuronal excitability through interactions with ion channels^2^. Polyamines have been linked to pain showing altered levels in human persistent pain states and modulation of pain behaviour in animal models^3^. However, the systems governing polyamine transport within the nervous system remain unclear. In undertaking a Genome Wide Association Study (GWAS) of chronic pain intensity in the UK-Biobank we found significant association with variants mapping to the *SLC45A4* gene locus. In the mouse nervous system SLC45A4 expression is enriched in all sensory neuron sub-types within the dorsal root ganglion including nociceptors. Cell-based assays show that SLC45A4 is a selective plasma membrane polyamine transporter, whilst the cryo-EM structure reveals a novel regulatory domain and basis for polyamine recognition. Mice lacking SLC45A4 show normal mechanosensitivity but reduced sensitivity to noxious heat and algogen induced tonic pain that is associated with reduced excitability of peptidergic nociceptors. Our findings thus establish a role for neuronal polyamine transport in pain perception and identify a new target for therapeutic intervention in pain treatment.

---

Chronic pain affects around one in five of the adult population and has a major negative impact on quality of life with profound socioeconomic consequences^4,5^. Indeed, when all pain conditions are taken, into account chronic pain is the leading cause of disability worldwide^6^. Unfortunately, current treatments are often inadequate due to poor efficacy and tolerability^5,7^. The molecular causes of chronic pain are not fully defined although enhanced excitability of nociceptors (those sensory neurons which detect injurious stimuli) and the central nociceptive circuits to which they project are important determinants of sensitised pain states^8,9^.

One group of endogenous metabolites that has been proposed to contribute to chronic pain are the polyamines which include putrescine (Put), spermine (Spm) and spermidine (Spd). These are ubiquitous polycationic alkylamines that have important roles in the synthesis, maintenance and stability of nucleic acids, cell signalling (including the stress response) and growth but can also modulate ion channel function^1^. Serum and tissue polyamine levels are raised in pain states such as inflammation and rheumatoid arthritis^10,11^. Injection of polyamines into the paw^3^ or spinal intrathecal space^12^ elicits pain behaviour (in rodents) and blocking their synthesis has been reported to reduce inflammatory pain^13^. However, it is unclear from previous studies as to the direction of effect, locus of action or how best to therapeutically intervene. The regulation of ion channels by polyamines is critically dependent on their location (intracellular versus extracellular) ^14,15^ and so understanding their transport is essential in determining the impact on neuronal function. Furthermore, excess polyamines are toxic and therefore tight cellular homeostasis is required to preserve cellular function but how this is achieved in the body is still unclear^16^.

To date only intracellular polyamine transport systems have been identified, which include the lysosomal polyamine transporter ATP13A2 (PARK9) ^17^ which is linked to Parkinson’s Disease and the vesicular polyamine transporters ATP13A3 and SLC18B1 (VPAT) ^18,19^ leaving the identity of the plasma membrane polyamine system unknown. Here we demonstrate that SLC45A4 encodes a plasma membrane polyamine transporter genetically linked to chronic pain in the human population, which presents an opportunity to understand the regulatory network linking polyamine biosynthesis and neuronal excitability.

## Variants in SLC45A4 are associated with altered pain perception in humans

Pain has a significant heritable component that varies depending on the sub-type of pain studied but median heritability in twin studies is 36%^20^. To investigate the genetic factors we undertook a genome-wide association study (GWAS) using data from the enhanced pain phenotyping questionnaires administered in the 2019 UK-Biobank (UKB) ^21^, enabling accurate case definition of chronic pain^22^. As the outcome measure, we used reported pain intensity on a numerical rating scale for the most bothersome chronic pain (n= 132,552, European ancestry, pain distribution shown in Fig. 1a). We identified a total of 29 genome-wide significant single nucleotide polymorphisms (SNPs) associated with pain intensity, including two independent loci with lead SNPs (illustrated in Fig. 1b and detailed in Extended data Table 1, and top 100 GWAS results in SI table 1): rs3905668 (p = 1.22 x 10^-8^) which is intergenic and near the *MSL2* (MSL Complex Subunit 2) gene and rs10625280 (p = 3.37 x 10^-8^) which maps to the *SLC45A4* (Solute Carrier Family 45 Member 4) gene.

**Figure 1.**
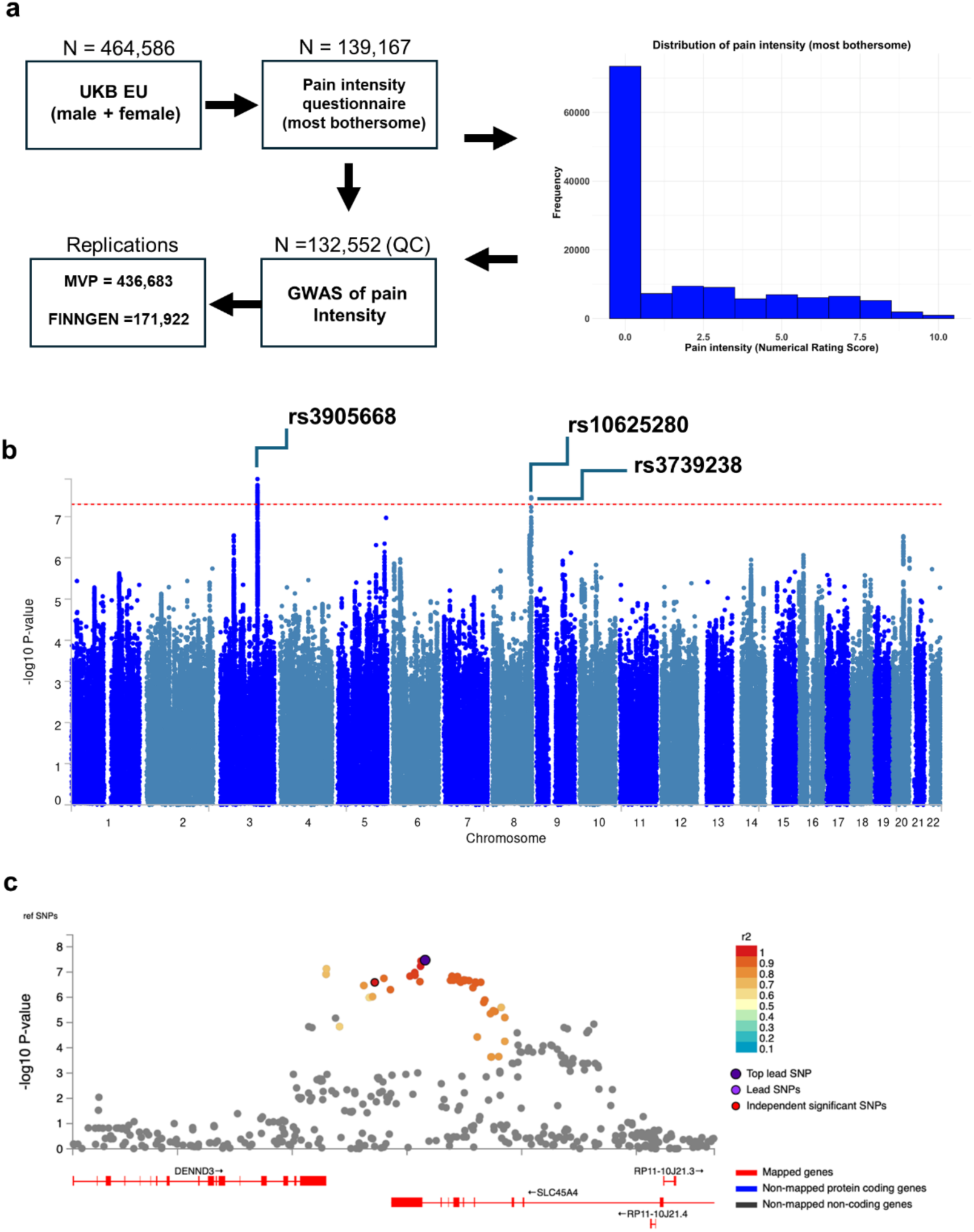
UKB pain intensity (most bothersome) GWAS identifies novel signals. **a,** A histogram illustrates the frequency distribution of pain intensity scores among participants, measured using the Numerical Rating Scale. Pain scores range from 0 (no pain) to 10 (worst possible pain). **b,** In a Manhattan plot from the UKB pain intensity GWAS, the genome-wide significant threshold is demarcated by a horizontal red line at 5 x 10^-8^. **c,** A regional plot centred on rs10625280, the principal SNP within the *SLC45A4* gene, elucidates associations within this genomic region. SNPs are differentiated by a colour gradient based on their linkage disequilibrium (r^2^ values) with the top independent significant SNP. The top lead SNPs in genomic risk loci, lead SNPs and independent significant SNPs are distinctly marked—encircled in black and highlighted in dark-purple, purple and red, respectively.

We focused on the *SLC45A4* locus (locus zoom plot Fig. 1c) given that it encodes a solute carrier (SLC) transporter, the function of which was unclear (although it had previously been proposed as a sucrose-proton transporter^23^). rs10625280 is situated within an intron of the *SLC45A4* gene (Fig.1c), with a subset of SNPs demonstrating linkage disequilibrium (R^2^ > 0.6), localising to the same genomic region. One of these is a missense variant in *SLC45A4* (r^2^ = 0.98, rs3739238, p.Asn718Asp, p = 3.63 x 10^-8^) which was significantly associated with pain intensity. Moreover, both SNPs were identified as an expression quantitative trait locus (eQTL) for *SLC45A4* and *DENND3* (DENN Domain Containing 3) across several databases including Open Targets Genetics, EyeGEx, GTEx, and eQTLGen database. The locus-to-gene (L2G) analysis pipeline prioritised *SLC45A4* with an overall L2G score of 0.86 (rs3739238) ^24^.

Having identified a genetic link between SLC45A4 and pain perception in the UKB-data, we then analysed data from the Million Veteran Program (MVP) in which a recent multi-ancestry genetic study of pain intensity in approximately 600,000 veterans identified 125 independent genetic loci, one of which was *SLC45A4* ^25^. The top SNP identified in MVP was rs59918340. The direction of association was dependent on ancestry. A review of the SNPs identified in our UKB GWAS of chronic pain intensity successfully replicated both the lead SNP rs10625280 (p = 1.21 x 10^-8^) and missense variant rs3739238 (p = 7.24 x 10^-9^) in the European ancestry GWAS of pain intensity in MVP (Extended data Table 1). The Finnish Genetics project (FinnGen) ^26^ GWAS also confirmed the association of these two variants with pain (Extended data Table 1). The direction of effect was consistent across different databases and studies, underscoring the robustness of these genetic associations.

We employed a Phenome-Wide Association Analysis (PheWAS) to examine secondary phenotypes linked with genetic variants rs10625280, rs3739238, and the *SLC45A4* gene, utilising data from the UKB, FinnGen, and the GWAS Catalog. rs10625280 showed significant association with 25 different traits, including skeletal (osteoarthritis) and several immunological traits concerning blood cell parameters (SI table 2 shows traits with significant association following Bonferroni correction). rs3739238 showed significant association with 35 traits, spanning immunology, skeletal (osteoarthritis), metabolism, psychiatry, and reproduction (SI table 2). The *SLC45A4* gene demonstrated associations with 55 distinct traits across a variety of health domains such as skeletal, reproductive, psychiatric, immunological, cardiovascular, neurological, and metabolic functions (SI table 2). Additionally, the Open Targets Genetics’ locus-to-gene pipeline^24^ identified a strong association between *SLC45A4* and multisite chronic pain (p = 2 x 10^-8^), highlighting its potential role in pain perception.

## SLC45A4 is a polyamine transporter

We next sought to understand the function of SLC45A4 at the molecular level. Based on distant homology to the plant sucrose transporter (SUC) in *Arabidopsis thaliana*, SLC45A4 was proposed as a proton coupled sucrose transporter ^23^. However, the human SLC45A4 protein only shares ∼ 26 % sequence identity to the *A. thaliana* SUC1 transporter, which coupled with recent studies reporting conflicting functions for this protein in metal ion transport^27^, leaves the role of SLC45A4 in the cell an ongoing enigma. We therefore sought to identify possible substrates using a correlation analysis between metabolomics and expression datasets, which can predict substrates for SLC proteins^28^. Our analysis highlighted that the expression of SLC45A4 across more than 1000 cell lines in the Cancer Cell Line Encyclopedia correlated positively to the levels of g-aminobutyric acid (GABA) (Extended Data Fig. 1a). However, neither GABA nor sucrose elicited a change in the thermal stability of purified SLC45A4 (Fig. 2a and Extended Data. Fig. 1b), prompting us to consider alternative metabolites. Although GABA is commonly synthesised from glutamate via glutamate decarboxylase enzymes^29^, an alternative pathway is available via the degradation of the polyamine Put^30^. We therefore tested a panel of substrates involved in GABA synthesis via the arginine/ornithine/putrescine (AOP) pathway. Surprisingly we observed a notable decrease in thermal stability in the presence of biogenic amines, with the polyamines Spm and Spd eliciting the largest response (Fig. 2a & Extended Data Fig. 1b). Cell-based radioactive uptake assays confirmed that SLC45A4 functions as a non-selective polyamine transporter with a pH optimum of between pH 7.5 – 8.0 (Fig. 2b, c) and no inhibition observed in the presence of L-lysine, L-ornithine or L-arginine (Extended Data Fig. 1c). Additionally, our data reveal that SLC45A4 is also selective in polyamine recognition. The longest polyamine, Spm, has an IC_50_ of 240 mM, followed by Spd (123 mM), with the smaller polyamines Put and cadervine (Cad) having the highest affinity (74 mM and 67 mM respectively) (Fig. 2d). Although it was long recognised that polyamines play a fundamental role in modulating neuronal excitability^31^, the identity of a specific neuronal biogenic amine transporter has remained mysterious. Our results clearly identify SLC45A4 as a plasma membrane polyamine transporter.

**Figure 2.**
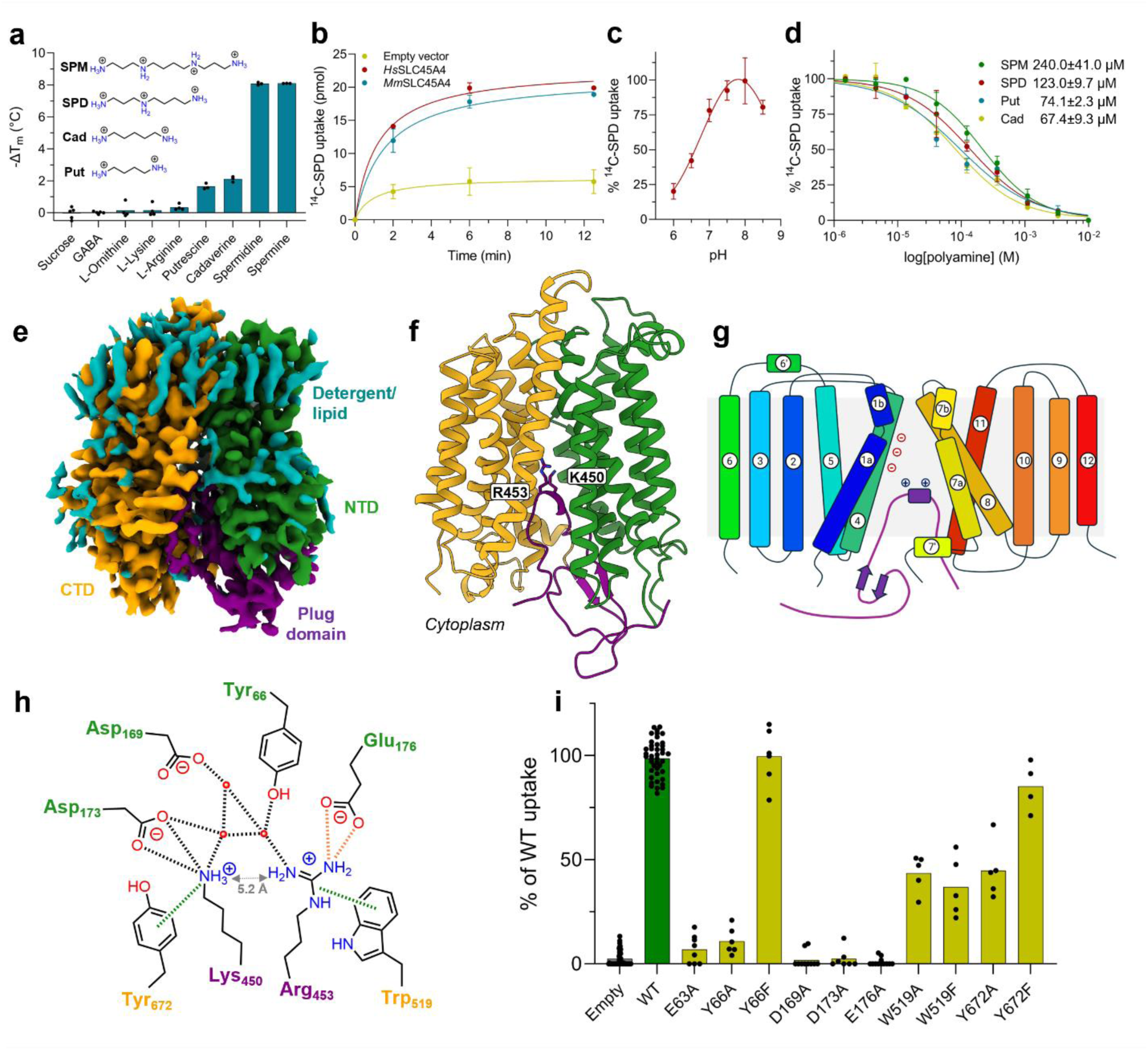
SLC45A4 is a polyamine transporter with a novel plug domain. **a**, Thermal stabilisation of SLC45A4 in the presence of metabolites in the AOP pathway using thermal stabilisation. n = 3 independent experiments, calculated mean and s.d. values are shown. **b**, Time course of ^14^C-SPD uptake in neuronal N2 cells overexpressing either human (*Hs*) or mouse (*Mm*) SLC45A4 in comparison to an empty vector control. n = 3 independent experiments, calculated mean and s.d. values are shown. **c**, Comparison of SLC45A4 activity in neuronal N2 cell under different external pH values. n = 3 independent experiments, calculated mean and s.d. values are shown. **d**, Polyamine competition of ^14^C-SPD uptake into neuronal N2 cells. The calculated mean (from 5 independent experiments) half-maximal inhibitory concentration values are shown ± s.d. **e**, Cryo-EM density of human SLC45A4 in LMNG:CHS detergent, contoured at a threshold level of 0.25. **f**, Cartoon representation of SLC45A4. The two 6TM bundles of the MFS fold are coloured by domain and the plug domain, which inserts in between the two is shown in purple. **g**, Topology diagram of SLC45A4, coloured blue to red from the amino terminus. **h**, Schematic of interactions between Lys450 and Arg453 in the plug domain and the polyamine binding site in SLC45A4. Hydrogen bonds and salt bridges are shown as black dashed bonds, cation-π bonds as green dashes and the charge interaction between Arg453 and Glu176, only observed in the nanodisc structure, as orange dashes. Residues are coloured by domain, as in panels e and f. **i**, ^14^C-SPD uptake in neuronal N2 cells overexpressing SLC45A4 and mutants of residues shown in **h**. n > 4 independent experiments.

## Cryo-EM structure of SLC45A4

To understand the structural basis for polyamine transport we proceeded to determine the structure of human SLC45A4 using cryo-EM. We determined the structure in both detergent and nanodisc to 2.80 and 3.25 Å respectively (Fig. 2e, SI Table 3 and Extended Data Fig. 2 & 3). The structures are similar with a root mean square deviation of 0.32 Å over 425 C_α_ atoms (Extended Data Fig. 4a). SLC45A4 consists of 12 trans-membrane spanning alpha helices which adopt the canonical Major Facilitator Superfamily (MFS) fold consisting of two six helical bundles in an inward open state^32^ (Fig. 2f, g). A unique feature of SLC45A4 is the presence of a large 25.5 kDa cytoplasmic domain inserted between TM6 and TM7. Although the majority of this domain is disordered in the volume, we observe clear density between Arg414 and Gln462, which extends into the canonical MFS binding site formed between the two six-helical bundles and plugs the transporter in an autoinhibited state by packing against the intracellular gating helices TMs 4 and 5 and TMs 10 and 11 (Fig. 2e and Extended Data Figs. 3e & 4a). The plug domain (414-462) forms an extended coil that places two conserved basic side chains, Lys450 and Arg453, into a solvated central cavity within the transporter domain that is noticeable for the extreme negatively charged surface, which facilitates the anchoring of the plug domain into the cavity (Extended Data Fig. 4b). Lysine 450 interacts with Asp173, which together with Asp169 and Glu176 form an acidic ladder on TM4 and, through a cation-pi interaction with Tyr672 (TM11) (Fig. 2h and Extended Data Fig. 4c). Arginine 453 interacts with Tyr66 (TM1) and makes a cation-pi interaction with Trp519 (TM7), which together with Tyr66 and Tyr672 is one of several conserved aromatic side chains within the binding site. Arg453 also interacts with a water molecule network linking the side chain to Asp169. Additional interactions between the plug domain and the transporter are also observed. Most notably, Ser452 hydrogen bonds with Glu176, while Tyr459 interacts with Arg180 (TM4), which serve to further stabilise the plug domain against the intracellular gating helices in the transporter. Finally, we noticed that in the nanodisc structure the membrane thins ∼ 9 Å where the plug domain enters the transporter (Extended Data Fig. 4d). Previous structures of the lysosomal polyamine transporter, ATP13A2, proposed a mechanism of membrane destabilisation to facilitate polyamine exit into the cytoplasm^33^. Our structure suggests a similar mechanism for polyamine release may operate within SLC45A4.

Efforts to capture a substrate bound state using both detergent and nanodisc samples however proved unsuccessful. However, like polyamines, both lysine and arginine contain primary and secondary amine groups. The positioning of these side chains within the canonical MFS binding site implies the plug domain functions as a pseudo substrate to auto inhibit the transporter. Interestingly, the distance between the amines in Lys450 and Arg453 side chains is ∼ 5 Å (Fig. 2h & Extended Data Fig. 4c), which is close to the length of the higher affinity substrates Put and Cad (Fig. 2a), suggesting that polyamines may interact similarly with the binding site. We therefore used the interactions made by Lys450 and Arg453 to guide our mutational analysis. The acidic ladder on TM4 is both strictly conserved within the SLC45A4 family (Extended Data Fig. 1d) and essential for polyamine uptake (Fig. 2i), with alanine variants showing no detectable transport in our cell-based assay. Also essential is Glu63 (TM1), which is another conserved acidic side chain that sits close to Glu176. In the nanodisc structure Arg453 adopts a different rotamer and interacts with Glu63 (Extended Data Fig. 4c), suggesting this side chain also interacts with polyamine substrates along with the side chains in the acidic ladder. However, whilst Tyr66Ala and Tyr672Ala mutants showed reduced activity, conservative mutations to phenylalanine showed wild-type (WT) levels of transport, suggesting bulky hydrophobic side chains at these positions facilitate transport. However, Trp519 was less important as mutating it to either alanine or phenylalanine only reduced function to around ∼ 50 % compared to WT levels. The types of interactions we observed within the binding site of SLC45A4 are consistent with those reported for the lysosomal polyamine pump ATP13A2 (PARK9) ^33^ and polyamine binding protein, PotD, from *Escherichia coli*^34^ (Extended Data Fig. 4e), suggesting that these two structurally distinct proteins use a combination of ionic and cation-pi interactions to recognise and transport polyamines across the membrane.

## SLC45A4 is highly expressed in sensory neurons

Little is known about the expression of SLC45A4 along the pain pathway, so we aimed to characterise its expression in neural tissues. Published single-cell RNAseq data-sets of the mouse nervous system shows that *SLC45A4* mRNA is broadly expressed but with a preponderance in dorsal root ganglia (DRG) sensory neuron subtypes^35–38^ (Extended data Fig. 5). Using qRT-PCR in WT C57Bl6 mice we confirmed that SLC45A4 expression was enriched in the DRG across the sensory neuraxis (Fig. 3a). *Insitu* hybridization of DRG sections showed that virtually 100% of DRG neurons expressed SLC45A4 (Fig. 3b) and it was expressed by all sub-types; peptidergic nociceptors (neuropeptide CGRP+), non-peptidergic nociceptors (isolectin B4 binding), C-low threshold mechanoreceptors (tyrosine hydroxylase+) and myelinated mechanoreceptors and proprioceptors (neurofilament heavy chain, NF200+) (Extended data Fig. 6) with higher expression in large diameter (NF200+) afferents. We also confirmed *SLC45A4* mRNA expression in the mouse spinal cord, with high ventral horn expression and limited dorsal horn (laminae III & IV) expression (Extended data Fig. 6). Existing data also show expression of SLC45A4 in human sensory neurons (Extended data Fig. 5). When electroporated into DRG neurons eGFP-tagged SLC45A4 is trafficked to the plasma membrane (Fig. 3c) consistent with a role in membrane transport.

**Figure 3.**
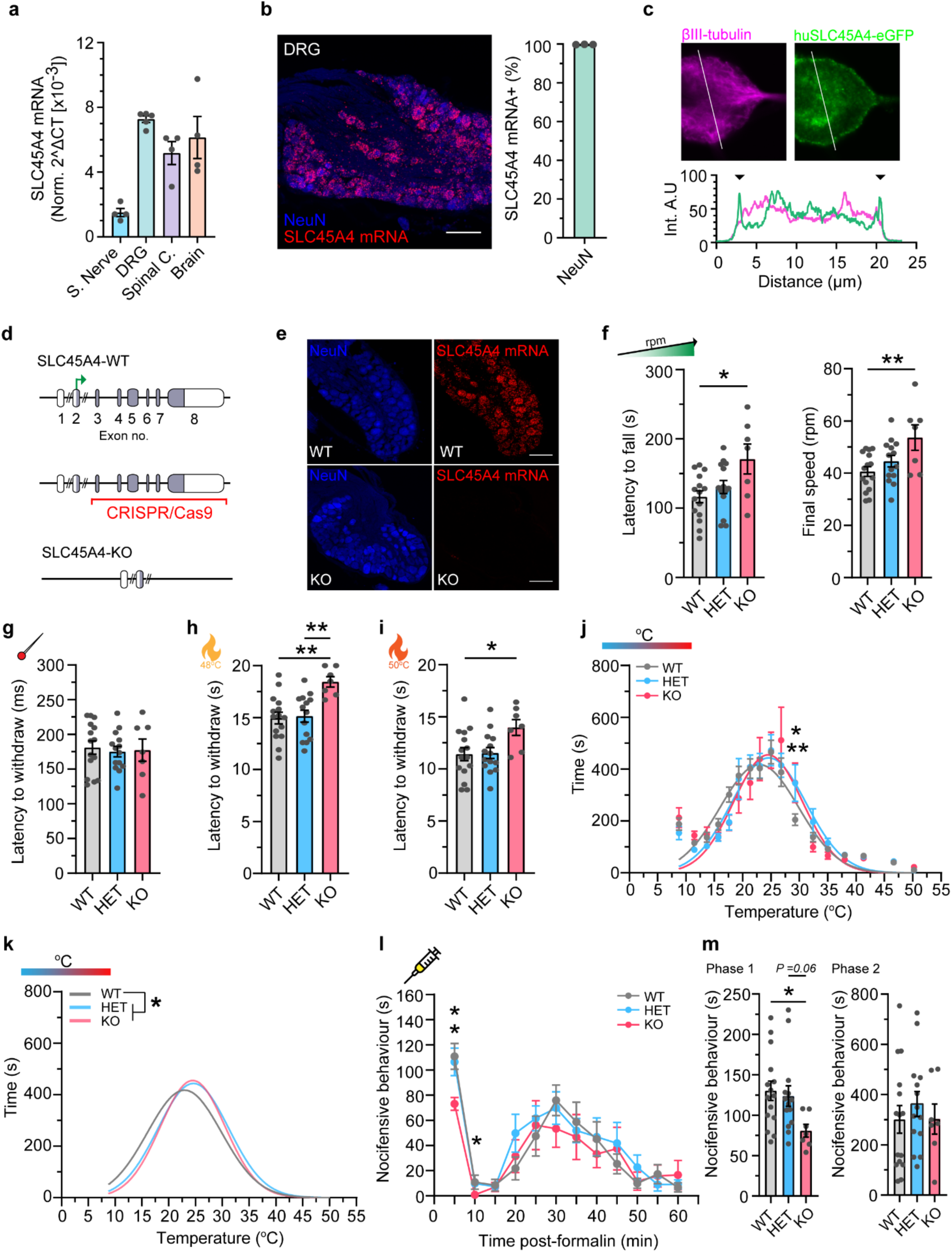
SLC45A4 is important for motor endurance, heat sensitivity and tonic pain. **a,** qPCR analysis of *SLC45A4* mRNA in tissues along the sensory neuraxis. From left to right, Sciatic nerve (n = 4 mice), Dorsal root ganglion (n = 5 mice), Spinal cord (n = 4 mice), Brain (n = 4 mice). **b,** *SLC45A4* mRNA is expressed in all NeuN positive mouse DRG neurons (n = 3 mice, 1777 cells). **c,** Human SLC45A4-eGFP transfected in to mouse sensory neurons localises to the plasma membrane. Black arrows in the profile plot indicate the increased intensity of huSLC45A4-eGFP at the membrane. **d,** Schematic illustrating the *SLC45A4* KO strategy using CRISPR/Cas9 deletion of exons 3-8. **e,** RNAscope ISH of *SLC45A4* mRNA in WT and KO DRGs. *SLC45A4* mRNA is absent in *SLC45A4* KO mice. **f,** *SLC45A4* KO mice show a longer latency to fall and a higher final speed, when challenged with a rotarod that gradually increases in speed (* P = 0.013, ** P = 0.0085). **g,** *SLC45A4* HET and KO mice show a normal latency to withdraw from a noxious pin-prick (P > 0.05). **h,** *SLC45A4* KO mice take longer to withdraw from a 48°C hotplate compared to WT and HET mice (** P = 0.002 and P = 0.0039 respectively, 20 s is the cut-off). **i,** *SLC45A4* KO mice have an increased latency to withdraw from a 50°C hotplate compared to WT mice (* P = 0.039, 20 s is the cut-off). **j,** Mice were given free roam of a thermal gradient apparatus where HET and KO mice had a rightward shift in their thermal gradient profile, towards warmer temperatures (two-way ANOVA, with post hoc Holm-Sidak’s test, WT vs HET ** P = 0.009, WT vs KO * P = 0.027). Data points and non-linear fit gaussian curves shown. **k,** The non-linear fitted gaussian curves for the thermal gradient profiles of WT, HET, KO mice. There is a rightward shit in the HET and KO curves. A single curve cannot explain all data sets, a different curve for one data set, WT. (Extra sum-of-squares F test, * P =0.024). **l,** *SLC45A4* KO mice display less nocifensive behaviours to a formalin injection compared to WT and HET mice (RM two-way ANOVA, post hoc Holm-Sidak’s test, 5 min: WT vs KO * P = 0.011, Het vs KO * P = 0.027. 10 min: HET vs KO * P = 0.018). **m,** The formalin test subclassed into Phase 1 (0-15 mins) and Phase 2 (15-60 mins). *SLC45A4* KO mice show less nocifensive behaviours in Phase 1 compared to WT mice (Kruskal-Wallis test, * P = 0.02). Phase 2 is normal in all groups. All data mean ± s.e.m. f-m: WT n = 15 mice, HET n = 14 mice, KO n = 7 mice. F-i and m: one way ANOVA, with Tukey post-hoc test. Scale bars 100 µm.

## Ablation of SLC45A4 impacts pain related behaviour and motor endurance

To explore the link between SLC45A4 function and pain perception we generated homozygote *SLC45A4-/-* knockout (KO) mice using CRISPR/Cas9 technology (Fig. 3d & Extended Data Fig. 7). Animals were born with no observable defects or perinatal mortality however we did note a lower than expected frequency of homozygote KO mice (SI Table 4), suggesting possible early embryonic lethality. Weight gain was normal (SI Table 4) however we noted a transient white hair phenotype resulting in a ‘salt and pepper’ appearance of the coat which appeared around day 20 and then faded and was absent by day 70 (Extended data Fig. 7). This finding is identical to that reported in an independent *SLC45A4* KO mouse (in which exon 5 was ablated) ^39^ and was proposed to be due to a transient defect in melanoblast differentiation; indeed Spd has been shown to enhance melanin production^40^. We undertook behavioural testing from the age of 10 weeks when hair colour had normalised, and the experiments could be blinded. No deficits were observed on open field behaviour tests for locomotion and exploration (Extended data Fig. 8 a-e). However, during a rota rod task of motor performance, where mice were challenged to run on a rotating rod that steadily increased in speed (rpm), the latency to fall from the rota rod and the maximum final speed were both increased in KO mice relative to WT (Fig. 3f), indicative of increased motor endurance. Anatomically, neuromuscular junctions in *SLC45A4* mutant mice were normal (Extended data fig. 9 a-b).

We next tested sensory behaviour. The reflex withdrawal responses of the mouse hindpaw were assessed in response to mechanical stimuli. The withdrawal threshold to a light touch stimulus (using von Frey hairs, Extended data Fig. 8F), or the latency to withdraw from a noxious pin prick stimulus (Fig. 3g), were unchanged in *SLC45A4* mutant mice. However, *SLC45A4* KO mice showed an increased latency to respond (i.e., hyposensitivity) to being placed on a 48 °C (Fig 3h) or 50 °C (Fig 3i) hot plate, compared to WT and Heterozygous mice. Sensitivity to a 53°C hotplate and a Hargreaves radiant heat source was unchanged (Extended data Fig 8g).

However, when assessing time spent in different regions of a thermal gradient apparatus we observed a notable difference in the distribution of the time spent in temperature zones, with KO and HET mice shifting their preference towards warmer temperatures compared to WT mice (Fig. 3 j & k). There was no difference in the reflex withdrawal latency to dry ice (noxious cold) (Extended data Fig. 8h). Our results demonstrate an altered thermal coding in *SLC45A4* mutant mice that appears specific for heat sensation over cold. Finally, we examined the nocifensive behaviours (paw lifting, licking and shaking) in response to intra-plantar administration of the algogen formalin, as a model of tonic pain. This behaviour occurs in two phases. The first phase is thought to primarily relate to direct activation of nociceptors through the ligand gated ion channel TRPA1^41,42^ and the second phase occurs due to spread of this activation and also spinal sensitisation. *SLC45A4* KO mice showed a significant reduction in their nocifensive behaviours during the first phase of the formalin response, compared to WT and HET mice (Fig. 3 l & m). As our expression studies and behavioural data both pointed towards a role of SLC45A4 in sensory neurons, we undertook anatomical and electrophysiological assessment of DRG physiology. There was no loss or change in sensory neuron subpopulation distributions (Fig. 4 a & b) and normal cutaneous nerve fibre density in *SLC45A4* KO mice compared to WT (Fig. 4 c & d), suggesting no major developmental phenotype in regard to sensory neuron survival or axon-outgrowth. On undertaking patch-clamp analysis of dissociated sensory neurons, we focused on two major nociceptor populations, non-peptidergic nociceptors (IB4-binding) and peptidergic nociceptors (defined in culture as small DRG neurons that do not bind the lectin IB4). We observed normal passive membrane properties (Extended data table 2) and comparable action potential thresholds (rheobase, Fig. 4 e & f) in *SLC45A4* KO neurons compared to WT. Interestingly, while non-peptidergic nociceptors were normal (Fig.4 g & h), we found a selective reduction in the supra-threshold excitability of peptidergic nociceptors from *SLC45A4* KO mice, in response to static (step) and dynamic (ramp) current injections, compared to neurons from WT mice (Fig. 4 i & j).

**Figure 4.**
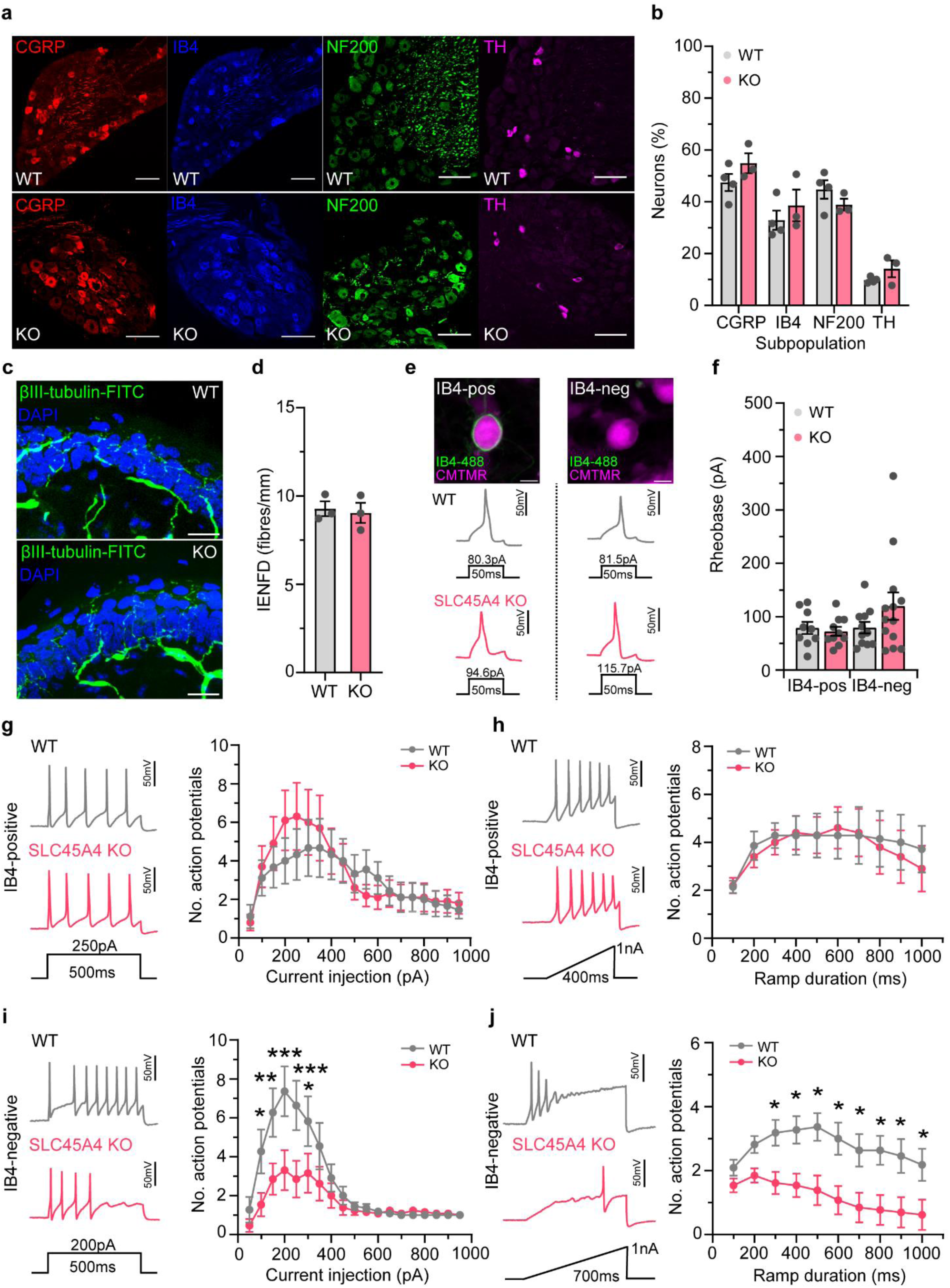
SLC45A4 regulates the excitability of peptidergic nociceptors. **a,** Examples of sensory neuron subpopulation markers in WT and KO mice. **b,** Quantification of each subpopulation marker as a percentage of total DRG neurons, highlights no developmental difference in DRG subpopulations between WT and KO mice. WT n = 4 mice (no. cells: 1073 CGRP, 747 IB4, 742 NF200, 167 TH), KO n = 3 mice (no. cells: 761 GCRP, 524 IB4, 402 NF200, 151 TH, two-way ANOVA with post-hoc Holm-Sidak test, P > 0.05.) Scale bars 100 µm. **c,** Example images of intra-epidermal nerve fibre density (IENFD) in WT and KO mice, scale bars 20 µm. **d,** Quantification of IENFD highlights no loss of nociceptive nerve terminals in glabrous skin of KO mice, compared to WT (WT n = 3 mice, 303 fibres, 9 sections, KO n = 3 mice, 312 fibres, 9 sections, unpaired t-test, P > 0.05). **e,** Example images of an IB4-positive and IB4-negative small sensory neuron, distinguished by the presence or absence of IB4-488 live binding, scale 10 µm. Bottom: example traces of action potentials elicited by a threshold current injection, from IB4-postive and IB4-negative neurons from WT and KO mice. **f,** Threshold excitability (rheobase – minimum current required to elicit and action potential) is similar between WT and KO small sensory neurons that are IB4-postive or IB4-negative (IB4-pos: WT n = 9 cells, KO n = 10 cells, IB4-neg: WT n = 11 cells, KO n = 13 cells, Kruskal-Wallis test with Dunn’s multiple comparisons test, P > 0.05). **g,** IB4-postive nociceptors from KO mice show normal repetitive firing patterns to step-current injections, compared to WT. (RM two-way ANOVA, post hoc Holm-Sidak’s test, P > 0.05). **h,** IB4-postive nociceptors from WT and KO mice are comparable when challenged with ramp-current injections (WT n = 7 cells, KO n = 10 cells, RM two-way ANOVA, post hoc Holm-Sidak’s test, P > 0.05)**. i,** IB4-negative nociceptors (peptidergic nociceptors) from KO mice are hypo-excitable to suprathreshold step-current injections, compared to WT neurons (RM two-way ANOVA, post hoc Holm-Sidak’s test, 100 pA * P = 0.039, 150 pA ** P = 0.0027, 200 pA *** P = 0.0002, 250 pA *** P = 0.0006, 300 pA * P = 0.047)**. j,** Peptidergic nociceptors from KO mice are hypo-excitable to ramp-current injections, compared to WT neurons (RM two-way ANOVA, post hoc Holm-Sidak’s test, 300-1000ms * P = 0.036, 0.021, 0.011, 0.013, 0.021, 0.015, 0.021, 0.036, respectively).

In summary, our results provide compelling evidence that *SLC45A4* is a neuronal membrane polyamine transporter that shows genetic association with human pain. It is expressed in sensory neurons and ablation of this function results in altered thermal coding and pain perception in mice. These behavioural changes are accompanied by a marked reduction in the excitability of peptidergic nociceptors which are known to encode thermal pain and the response to many algogens. Polyamines are known to functionally modulate a number of ion channels that may be candidates for impacting on excitability of nociceptors including: the Transient Receptor Potential family (TRPV1, 3,4) ^14,15^, inward rectifier K^+43^ and acid-sensitive ion channels (ASICs) ^44^. Our study has identified *SLC45A4* as the much anticipated neuronal polyamine transporter and suggests that *SLC45A4* may be a promising molecular target to modulate pain perception in humans.

## Methods

### Study Population

The UK Biobank (UKB) is a prospective cohort study that has collected extensive phenotypic and genetic data from approximately 500,000 participants across the United Kingdom^1,2^. All participants provided written informed consent.

### Phenotype description and assessment

Phenotypes were defined using the “Experience of Pain” UK biobank self-assessment questionnaire data under project ID 49572. This is part of the online UKB follow-up and was sent in May 2019 to all UK Biobank participants, with an active email address, who had consented to further electronic contact (n = 335,587). Definitions and characteristics of the phenotypic endpoints are described in detail in Baskozos et al., 2023^3^. Participants who have withdrawn (153 as of 15/09/2023) were excluded from the analysis. Chronic pain was defined by considering a screening question asking whether participants are “troubled by pain or discomfort present for more than 3 months” (item f120019). Question “area most bothered by pain in the last three months” (item f120037) was used to define the location of most bothersome pain and the intensity of the most bothersome pain was defined from items 120023 -120035 asking for the “rating of pain” in the most bothersome pain location. People with No chronic pain had their intensity values imputed to 0. People who self-reported fibromyalgia (f120009), chronic fatigue syndrome/myalgic-encephalomyelitis (f120010) or chronic pain all over the body (f120021) were excluded from the analysis (11,951 with chronic pain and 821 with no chronic pain). We had valid most bothersome pain ratings for 139,167 (71,904 and 67,263 meeting criteria for no chronic pain and chronic pain respectively).

### Genotyping, imputation, and quality control

In this study, we utilised genotype datasets from the UKB to investigate genetic factors related to chronic pain. We used two specific releases: version 2 for directly genotyped variants and version 3 for imputed genotypes. Initially, approximately 50,000 UKB participants were genotyped using the Applied Biosystems UK BiLEVE Axiom Array by Affymetrix. Subsequently, around 440,000 participants were genotyped with the Applied Biosystems UK Biobank Axiom Array.

Quality control and imputation procedures, as detailed previously^2^ and resulted in the released dataset of 488,377 samples and 805,426 directly typed markers from both arrays. This dataset was further quality controlled before phasing and imputation using a combined Haplotype Reference Consortium (HRC) and UK10K reference panel. The imputed dataset has over 90 million autosomal SNPs, short indels and large structural variants for 487,442 individuals.

To assess genetic ancestry, we employed the KING software^4^ (version 2.3.2), using the 1000 Genomes Project as a reference panel. The directly genotyped dataset was used with additional quality control filters using PLINK^5^ (version 1.90b6.21, https://www.cog-genomics.org/plink/1.9/) that included: autosomes, minor allele frequency ≥ 5%, not present in high LD regions and LD pruning using a R^2^ threshold of 0.2 with a window size of 50 markers and a step size of 5 markers. This analysis enabled us to identify five distinct subpopulations within the UK Biobank: African (9,059 samples), Ad Mixed American (605 samples), East Asian (2,572 samples), European (464,586 samples), and South Asian (9,604 samples). Additionally, there were 1,951 samples for which ancestry could not be determined and were thus categorized as missing.

Genotyping data, encompassing both directly genotyped and imputed variants, and excluding individuals who had withdrawn their consent from the UKB study were available for 487,071 samples. Additional quality control measures were applied that further excluded 367 samples where the reported sex did not match the inferred sex from their genetic data, 651 samples with suspected sex chromosome aneuploidy, and 188 samples with more than ten putative third-degree relatives. In total, 1,024 samples were excluded, with some samples falling into multiple exclusion categories.

Samples of European ancestry was selected, resulting in 462,402 samples available for analyses. The imputed dataset (chromosome 1-22) was restricted to common and low frequency variants (minor allele frequency ≥ 1%)) that were imputed with high confidence (imputation accuracy Info score ≥ 0.8) leaving 9,572,556 variants in the dataset.

### Association analyses and candidate SNPs identification

We conducted association analyses using REGENIE (version 3.4.1) ^6^. REGENIE employs a two-step whole-genome regression method that effectively accounts for population stratification and sample relatedness. For the continuous outcome, we applied rank-based inverse normal transformation to improve the distribution and meet the assumptions of the regression model. The model included the following covariates: age at the time of completing the questionnaire, sex, genotyping array, and the top 10 principal components provided by the UK Biobank. REGENIE step 1 was run on a set of the directly genotyped variants, filtered using PLINK2^5^ (version 2.00a5, https://www.cog-genomics.org/plink/2.0/) that included: sample genotyping rate ≥ 90%, autosomes, minor allele frequency ≥ 1%, Hardy–Weinberg equilibrium test not exceeding P = 1 × 10^−15^, variant genotyping rate ≥ 99%, not present in high LD regions and LD pruning using a R^2^ threshold of 0.9 with a window size of 1,000 markers and a step size of 100 markers. REGENIE step 2 was run on the imputed dataset.

To identify risk loci and their lead variants, we performed LD clumping using the Functional Mapping and Annotation of Genome-Wide Association Studies (FUMA) ^7^. We set a range of 250 kb and an r^2^ threshold of >0.6 to define independent significant SNPs. For lead SNPs, we used an r^2^ threshold of >0.1. The analysis was based on the respective ancestry from the 1000 Genomes Phase 3 EUR reference panel^8^. After clumping, we combined genomic risk loci within 1 MB of each other into a single locus. Additionally, we leveraged resources from Open Targets Genetics, which integrate data from human GWAS and functional genomics, including gene expression, protein levels, and chromatin interactions across various cell types and tissues^9^. This comprehensive approach enabled us to confirm the connections between GWAS-associated loci, variants, and their probable causal genes.

### Phenome-Wide Association Analysis (PheWAS)

To determine potential associations between the lead SNP associations and their corresponding genes from our study and additional traits, we conducted a Phenome-Wide Association Analysis (PheWAS). This analysis involved generating PheWAS plots from a comprehensive dataset comprising 4,756 GWAS summary statistics, acquired from the GWAS ATLAS. We incorporated all relevant GWAS and associated genes into our selection criteria. For the PheWAS SNP plots, SNPs were deemed significant with P values less than 0.05 and applied the Bonferroni correction method to adjust for multiple comparisons.

### Gene-Based Test, Pathway, and Enrichment Analyses

In our study, we used the FUMA software for gene-based tests, pathway exploration, and enrichment analyses. FUMA leverages GWAS summary statistics to prioritize genes, assess gene expression, and enrich pathway processes. To address the issue of multiple testing, FUMA applied the Bonferroni correction with a threshold of P_bon_ < 0.05. Additionally, FUMA incorporates multimarker analysis of genomic annotation (MAGMA) for both gene-based and gene-set analysis.

### Ethics

Data for this study were obtained from the UK Biobank for project “Risk factors for chronic pain,” Application ID: 49572. UK Biobank has approval from the North West Multicentre Research Ethics Committee (MREC) as a Research Tissue Bank (RTB) approval, REC reference: 21/NW/0157, IRAS project ID: 299116. This approval means that researchers do not require separate ethical clearance and can operate under the RTB approval.

### Metabolome Correlation analysis

Correlation analysis to identify potential substrates was performed as described^13^. Briefly, RNAseq raw count data for cell lines in the CCLE was downloaded and processed using median of ratios normalization. Metabolomics data was downloaded from the CCLE website (https://sites.broadinstitute.org/ccle/datasets). Cell lines were matched, and Spearman’s rank correlation coefficients were calculated between each SLC protein and each metabolite. Significance of the correlation coefficients were adjusted using the Benjamani Hochberg multiple testing method and plotted as a volcano plot with the value of the correlation coefficient on the x axis and the log10 of the significance on the y axis.

### Cloning, expression and purification of *Hs*SLC45A4

The gene encoding full-length human SLC45A4 (Uniprot Q5BKX6) was inserted into pDDGFP-Leu2D^14^, containing a C-terminal tobacco etch virus (TEV) protease cleavable eGFP-His_8_ tag, for expression in *S. cerevisiae* strain BJ5460 (ATCC-208285). Transformed yeast were cultivated in synthetic complete media without leucine (-Leu), supplemented with 2% (wt/v) glucose, at 30°C to an OD_600_ of 5.0-6.0 and diluted ninefold into -Leu with 2% (v/v) lactate pH 5.1. Once OD_600_ reached 1.8-2.2, expression was induced by addition of 1.5% galactose and expression maintained for 20-24 hr. Cells were harvested, lysed through high pressure cell disruption (40kpsi) and membranes isolated through ultracentrifugation at 200k g for 90 minutes. Membranes were washed in wash buffer, 20 mM HEPES pH 7.4, 1M KAc, and isolated again at 200k g for 90 minutes before being resuspended in 1xPBS and stored at -80°C.

For expression in tissue culture, SLC45A4 was inserted into the pLexM vector^15^ with a C-terminal Avi tag (GLNDIFEAQKIEWHE), followed by TEV-cleavable eGFP-His_6_ fusions. HEK293F cells were cultured in FreeStyle™ 293 Expression Medium (ThermoFisher Scientific) at 37°C and 8% CO_2_ and were, 24 prior to transfection, passaged to 0.7x10^6^ cells mL^-1^ to give a density of 1.3-1.4⋅10^6^ cells mL^-1^ upon transfection. Transfection was carried out with 1.1 mg plasmid and 2.2 mg linear polyethyleneimine (PEI) MAX (Mw 40,000; Polysciences Inc.) per L culture. Following transfection, sodium butyrate was added to 10 mM to increase protein expression. Cells were harvested 40 hr post-transfection. Membranes were prepared by lysing the cells via sonication and unbroken cells and cell debris were pelleted at 10,000g for 10 mins at 4 °C and membranes were harvested through centrifugation at 200,000g for one hour and washed once with 20 mM HEPES pH 7.5, 20 mM KCl. After washing the membranes were resuspended in PBS and snap frozen for storage at -80 until required. For purification, thawed membranes (∼ 10g wet weight) were solubilized in 130 ml of buffer containing 1 x PBS pH 7.4, 150 mM NaCl, 10% (v/v) glycerol and 2% (wt/v) *n*-dodecyl-β-D-maltopyranoside (DDM) with 0.4% (wt/v) cholesteryl hemisuccinate (CHS) for nanodisc reconstitution or 1% lauryl maltoside neopentyl glycol with (LMNG) 0.1 % (wt/v) CHS for 90 minutes at 4 °C under gentle agitation using a magnetic stir plate. Insoluble material was removed through centrifugation for one hour at 200,000g. SLC45A4 was purified to homogeneity using standard immobilised affinity chromatography (IMAC) protocols in *n*-either dodecyl-β-d-maltopyranoside (DDM) detergent (Anatrace) with cholesterol hemisuccinate (5:1 ratio DDM: CHS) or lauryl maltoside neopentyl glycol with (LMNG) 0.1 % (wt/v) CHS. In brief, 4mL of nickel NTA resin (Fisher Scientific) was added with 25mM imidazole (Sigma) for three hours at 4 °C under gentle agitation using a magnetic stir plate. The resin was loaded onto a gravity flow column (BioRad) and washed with eight column volumes (CVs) of buffer containing either 0.15 % DDM: CHS (5:1 ratio) or 0.15 % LMNG: CHS (5:1 ratio) and 25mM imidazole, followed by 15 CVs of buffer with 30 mM imidazole. The protein was eluted in four CVs of buffer containing 250 mM imidazole and dialysed overnight with TEV protease (1:0.5 M ratio) against 20 mM Tris pH 7.5, 150 mM NaCl with 0.02 % DDM and 0.004 % CHS or 0.003% LMNG: CHS (10:1 ratio) at 4 °C under gentle agitation using a magnetic stir plate. Following TEV cleavage, the protease and cleaved his tagged GFP were removed through nickel affinity chromatography as described above adding 10 mM imidazole to the dialysed sample. The protein was then concentrated to 500 μL using a 50KDa MWCO spin concentrator (Sartorius) at 4 °C and subjected to size exclusion chromatography (Superdex 200) in the same buffer as above.

### Cell-based ^14^C-SPD transport assays

Spermidine uptake activity of overexpressed SLC45A4 was assayed in Neuro-2A cells (ATCC CCL-131), using ^14^C-radiolabelled SPD (American Radiolabeled Chemical, Cat# ARC3138, 100 mCi/mmol). Cells were maintained in Gibco DMEM (high glucose, GlutaMAX Supplement, pyruvate, Cat# 31966021) under 37°C and 5% CO_2._ For transport assays, 1.0-1.7⋅10^5^ cells were seeded per well in 12-well plates and transfected 48-52 hr later (once confluency had reached ∼60-80%) using FuGENE HD (Promega Cat# E2311) transfection reagent, with 1 µg plasmid (wild-type SLC45A4 or mutants in pLexM, with a C-terminal FLAG tag) and 2.5 µL FuGENE per well. Fresh media was placed onto the cells 12-15 hr thereafter and assays carried out 40-46 hr following transfection.

Prior to measuring transport activity, cells were washed twice in assay buffer (25 mM HEPES pH 7.5, 135 mM NaCl, 5 mM KCl, 1.2 mM MgCl_2_, 28 mM glucose) and incubated for 3 min at 37°C following the second wash. Immediately after the incubation, 240 µL of the assay buffer supplemented with 1 µM ^14^C-SPD and 9 µM cold SPD were pipetted onto the cells and incubated for 2, 8 and 14 min at 37°C. Cells were then washed twice in 500 µL ice-cold assay buffer and the cells lysed in 20 mM Tris pH 7.5, 0.2% Triton X-100. The amount of transported ^14^C-SPD was calculated via scintillation counting in Ultima Gold (PerkinElmer) using a standard curve for the substrate. For assaying SLC45A4 variants, single timepoints at 8 min were measured. For competition assays and IC_50_ measurements, only 1 µM of ^14^C-SPD was used in the substrate mixture along with the desired concentration of the competing compound. Experiments were performed a minimum of six times to generate an overall mean and s.d. Expression of wild-type SLC45A4 and mutants were assessed using western blotting on membrane fractions with an anti-FLAG antibody (Merck F1804) at 5,000x dilution, using and anti-β-actin antibody (Merck A2228) (Extended data Fig 1e). Plasma membrane localisation was measured using immunofluorescence (Extended data Fig 1f). Cells were seeded on glass coverslips, transfected as described above with FLAG-tagged SLC45A4 and 36 hr post-transfection, cells were washed in PBS pH 7.4 and fixed in 4% paraformaldehyde for 7 min. Following quenching in 50 mM ammonium chloride and further washing, the cells were permeabilised in 100 µM digitonin and coverslips blocked in 1% bovine serum albumin (BSA) for 30 min. Cells were stained with mouse anti-FLAG (1:200 dilution) and rabbit anti- Na^+^/K^+^ ATPase (1:50 dilution) primary antibodies for 1 hr, washed and further stained with goat anti-mouse IgG AlexaFluor-488 (1:200) and anti-rabbit IgG AlexaFluor-647 (1:200) secondary antibody-fluorophore conjugates for imaging. Imaging was carried out on a LSM-980 confocal microscope (Zeiss) and images processed in ZenBlue (v3.9, Zeiss) and ImageJ^16^.

### Thermal stability measurements

For assessment of thermal stability in the absence or presence of compounds, nano differential scanning fluorimetry (NanoDSF) measurements were carried out on a Prometheus NT.28 instrument (NanoTemper Techonlogies). Purified SLC45A4 was diluted to a final concentration of 0.2-0.4 mg/mL in buffer (20 mM Tris pH 7.5/20 mM MES pH 7.5, 150 mM NaCl, DDM: CHS (0.03: 0.003%) with 2.5-50 mM metabolite. Unfolding was monitored as the ratio of Trp fluorescence emission at 330 nm and 350 nm between 20-90°C using a ramp rate of 1°C min^-1^. The apparent T_m_ was determined as the maximum of the 1^st^ derivative of the emission ratio.

### Nanodisc reconstitution

SLC45A4, purified from HEK293F in DDM:CHS, was reconstituted into MSP1D1 lipid nanodiscs with EBC lipid (85% (mol/mol) *E. coli* polar lipid extract, 10% bovine brain polar lipid extract, 5% cholesterol) using the BioBead method^17^. 100 µg SLC45A4 was mixed with MSP1D1 and EBC lipid solubilised in 0.5 M sodium cholate, at a molar ratio of 1:5:75, respectively, the mixture incubated on ice for 1 hr and excess detergent removed through stepwise addition of BioBeads and overnight incubation under gentle agitation. Insoluble material was removed through ultracentrifugation and reconstituted SLC45A4 separated from empty nanodiscs and excess lipid on a Superdex200 Increase 10/300 GL column in 20 mM HEPES pH 7.5, 150 mM NaCl and concentrated to 2 mg/mL for CryoEM analysis.

### CryoEM sample preparation and data acquisition

For the LMNG sample, purified SLC45A4 was subjected to a further round of SEC polishing on a Superdex200 Increase 10/300 GL column in 20 mM Tris pH 7.5, 150 mM NaCl, 0.001% LMNG:CHS (10:1), to separate from empty detergent micelles, and the unconcentrated peak fraction (0.31 mg mL^-1^) used for grid preparation. Sample was adsorbed to glow-discharged holey carbon-coated grids (Quantifoil 300 mesh, Au R1.2/1.3) for 10 s. Grids were then blotted for 2 s at 100% humidity at 10 °C and frozen in liquid ethane using a Vitrobot Mark IV (Thermo Fisher Scientific). Data were collected in counted mode in Electron Event Representation (EER) format on a CFEG-equipped Titan Krios G4 (Thermo Fisher Scientific) operating at 300 kV with a Selectris X imaging filter (Thermo Fisher Scientific) with slit width of 10 e-V at 165,000x magnification on a Falcon 4i direct detection camera (Thermo Fisher Scientific) corresponding to a calibrated pixel size of 0.732 Å. Movies were collected at a total dose of 57.6 e-/Å2 fractionated to ∼ 1 e-/Å2 per fraction. SLC45A4 reconstituted into EBC:MSP1D1 nanodiscs, as described above, was concentrated to 2 mg mL^-1^ and directly used for preparation of grids. 3 µL of sample were applied on a glow-discharged holey carbon grids (Quantifoil Cu R1.2/1.3 300 mesh), blotted for 5 s at 100% humidity and 4°C and plunge-frozen in liquid ethane using a Vitrobot Mark IV. Data were collected in counted super resolution bin2 mode on a Titan Krios G3 (FEI) with a K3 camera (Gatan) and BioQuantum imaging filter at 300 kV, with a pixel size of 0.832 Å. A total of 20,184 micrographs were collected using a dose of 15.89 e^-^/Å^2^/s and an exposure time of 2.5 s, giving a total dose of 39.71 e^-^/Å^2^.

### Cryo-EM data processing

Patched (20 x 20) motion correction, CTF parameter estimation, particle picking, extraction, and initial 2D classification were performed in SIMPLE 3.0^18^. All further processing was carried out in cryoSPARC 3.3.1^19^ and RELION 3.1^20^, using the csparc2star.py script within UCSF pyem^21^ to convert between formats. Global resolution was estimated from gold-standard Fourier shell correlations (FSCs) using the 0.143 criterion and local resolution estimation was calculated within cryoSPARC.

The cryo-EM processing workflow for SLC45A4 in LMNGCHS is outlined in Extended Data Fig. 2. Briefly, particles were subjected to two rounds of reference-free 2D classification (k=300 each) using a 140 Å soft circular mask within cryoSPARC. Four volumes were generated from a 479,080 particle subset of the 2D-cleaned particles after multi-class ab initio reconstruction using a maximum resolution cutoff of 6 Å. Particles from the most populated and structured class were selected and another multi-class ab initio reconstruction (k=4) performed. Output volumes were lowpass-filtered to 7 Å and used as references for a 4-class heterogeneous refinement against the full 2D-cleaned particle set (2,815,141 particles). Particles (964,648) from the most populated and structured class were selected and non-uniform refined against their corresponding volume lowpass-filtered to 15 Å, generating a 3.3 Å map. A multi-class ab initio reconstruction (k=4) using a maximum resolution cutoff of 6 Å was performed on these particles, generating four volumes that were lowpass-filtered to 7 Å and used as references for heterogeneous refinement against the same particle set. Particles (700,436) belonging to the two most populated and structured volumes (which were in opposite hands) were combined and subjected to non-uniform refinement against one of the corresponding volumes lowpass-filtered to 15 Å, generating a 3.3 Å map with slightly improved density over the previously refined volume. Bayesian polishing followed by non-uniform refinement (15 Å lowpass-filtered reference) further improved map quality to 3.0 Å. Per-particle defocus refinement followed by non-uniform refinement (15 Å lowpass-filtered reference) resulted in a 2.8 Å map that was used for model building.

For the SLC45A4 nanodisc structure (Extended Data Fig. 3), the 20,196,090 extracted particles (box size of 240 pix) were subjected to three rounds of conservative 2D classification, to give 1,473,487 particles which were further classified into four classes in *ab-initio* reconstruction. Three rounds of less conservative 2D classification of the initial particle set gave 5,812,766 particles which were classified by two rounds of heterogeneous refinement, using the best *ab-initio* map from the earlier reconstruction. After further sorting particles using *ab-initio* reconstruction, Bayesian polishing was carried out and particles re-extracted with a box size of 320 pix. After 2D classification, two further rounds of *ab-initio* reconstruction, non-uniform and local refinement gave a final map of 3.25 Å (where FSC=0.143) from 227,752 particles.

Model building was carried out in *Coot*^22^ (v0.9.8.1 EL) and ISOLDE^23^, refinement in PHENIX (v1.20.1-4487) real-space refinement^24^ and validation in MolProbity^25^. Images were generated using PyMol^26^ and ChimeraX^27^.

### Animals

Animals were group-housed in temperature and humidity controlled, specific pathogen-free facilities in individually ventilated cages. Mice were on a 12:12hr light-dark cycle with food and water provided ad libitum. All experiments were carried out using male and female C57BL/6J mice. Behavioural studies were carried out on age and sex matched littermates. All transgenic mice were backcrossed onto a C57BL/6J background. All procedures complied with the UK Animals (Scientific Procedures) Act (1986) and were performed under a UK Home Office Project Licence in accordance with University of Oxford Policy on the Use of Animals in Scientific Research. This study conforms to the NC3Rs policy on reduction, refinement and replacement of animal research, and to the ARRIVE guidelines^10^.

### Generation of the SLC45A4-KO mouse

Design and development of the SLC45A4-KO mouse was carried out in partnership with Taconic Biosciences and Cyagen Biosciences. Briefly, CRISPR/Cas9 technology was used to delete exons 3-8 of the mouse SLC45A4 gene on chromosome 15. This 13.3kb deletion accounts for 89.64% of the SLC45A4 coding region. Guide RNAs were designed to target only the SLC45A4 locus and injected together with Cas9 into mouse zygotes. Chimeric F0 offspring mice were bred with wildtype mice to generate heterozygous founders F1 which were screened for successful SLC45A4 editing. The targeted region of SLC45A4 was sequenced to confirm deletion of exons 3-8. Mice were genotyped using primers in SI table 5 using a standard Taq polymerase based protocol. WT band: 652bp, KO band: 468bp.

### Mouse Histology

#### Tissue collection

For histology studies, mice received an overdose of pentobarbital, then vascular perfused with saline and 4% paraformaldehyde (PFA). DRG, EDL muscle and skin tissue was collected and post-fixed in 4% PFA for 1-2h, Spinal cord for 24hr. DRG, skin and spinal cord were cryoprotected in 30% sucrose and stored for at least 48h before embedding in OCT. DRG tissue was cryo-sectioned at 12 μm, skin at 30 µm and spinal cord at 20 µm. Fixed EDL muscle was teased into small fibre bundles of about 1mm diameter per bundle. For histology on neural cultures, live CellTracker^TM^ dye CMTMR was added along with IB4-conjugated Alexa 488, or cells were washed and fixed in 4% PFA for 15 min RT.

#### Immunohistochemistry (IHC)

Standard immunohistochemistry techniques were used. Antigen retrieval in a citric acid buffer was performed on skin sections prior to IHC (Citric acid monohydrate, tween-20, dH_2_0, pH 6, 65oC, 15 min). Briefly, the sectioned or teased samples were washed in PBS and blocked in a blocking solution (5% normal goat serum and 0.3% TritonX-100, PBS) for at least 30 mins at room temperature (RT). Primary antibodies (SI table 6) were diluted in the same blocking solution and were left to incubate on the tissue samples overnight at RT or 4°C. Samples were washed in a washing buffer solution (0.3% TritonX-100, PBS). Secondary antibodies (SI table 6) were diluted in the washing buffer solution and left to incubate on the tissue for 1-2h at RT in dark conditions. Samples were washed thoroughly in the washing buffer, DRG and skin sections were mounted using Vectorshield (with or without DAPI), muscle fibres were mounted onto microscope slides (Avantor VWR, Cat. No 631-0108) using hard-set confocal matrix (Micro tech Lab). Images were taken on a confocal microscopes (Zeiss LSM-710, LSM-780 or Olympus FV3000).

#### In situ hybridisation (ISH)

In situ hybridisation (ISH) was performed as detailed in the user instructions for the RNAscope2.5 RED Chromogenic assay kit (Advanced Cell Diagnostics, Bio-techne, United Kingdom). Briefly, fixed DRG tissue underwent pretreatment with H_2_O_2_, protease III, and 100% ethanol. The tissue was then incubated at 2h at 40°C with an SLC45A4 mRNA probe (Cat. No. 522131). Probe amplification, washing steps and chromogenic development with fast red were carried out as detailed in the user kit. The tissue was then co-stained and imaged using the IHC protocol detailed above.

#### Image analysis

All image analysis was carried out using ImageJ/Fuji (NIH). For IHC, At least, three sections per animal were used from three animals per group. Total cells and cells positive for each subpopulation marker were counted using the multi-point tool. For ISH, Cells were defined and mRNA intensity calculated. Background intensity was calculated tissue that underwent the ISH protocol with a standard negative control mRNA probe. A threshold for positive cells was defined as cells whose average intensity was more than mean + 3*SD of background intensity. Cell size classification was as follows: cell area < 490μm^2^ was classified as a small cell, cell area between 490-962μm^2^ was classified as a medium cell and a large cell had an area greater than 962μm^2^. Intra-epidermal nerve fibre density (IENFD) analysis; nerve fibres crossing into the epidermis were counted live under the ocular lens of the Olympus FV3000. The distance of each section was calculated post-hoc in Fuji and nerve fibres quantified as fibres/mm. Spinal neurons were first segmented using Cellpose 2.0. Within Cellpose 2.0, we used a human-in-the-loop pipeline to train a custom model by annotating cells on the Cellpose graphical user interface (GUI). Cells were determined to be positive for target probe mRNA if signal intensity was 3*SDs above negative control readings. Superficial lamina (I+II) were defined by IB4 staining to mark lamina II. A reference template (Atlas of the Mouse Spinal Cord) was used analyse all other laminae. For assessment of neuromuscular junction (NMJ) size, fragmentation, area, and nerve terminal registration, optimised settings for each image were used. Between three and five Z-stacks were obtained per muscle sample. Ten NMJs were analysed from each animal using ImageJ software. The ImageJ macro (BetterAreaGreyValues) used to measure NMJ area and fragmentation (SI methods). Registration of different fluorescence signals was analysed using the JACoP plugin for Image J. All comparative image analysis was conducted while the experimenter was blinded to genotype.

### Quantitative real time PCR (qPCR)

Tissue was collected from mice following overdose with pentobarbital and trans-cardiac perfusion with ice cold saline. Tissues were rapidly dissected and flash frozen in liquid nitrogen and stored at -80°C. RNA was isolated from fresh-frozen mouse tissues using a combination of TriPure (Roche) and a RNeasy mini kit (Qiagen). Briefly, tissue was homogenised in Tripure using a handheld homogeniser (Cole-Parmer), treated with chloroform and then column-purified and eluted in RNase-free water. Genomic DNA was removed with on-column DNase I digestion. cDNA synthesis was carried out using EvoScript Universal cDNA Master kit (Roche). Gene expression was quantified by detecting amplified material using LightCycler SYBR Green Master Mix (Roche) on the LightCycler 480 II system (Roche). Three technical replicates were included for each sample. Results were normalized to three reference gene controls (mouse Actb, Gapdh, and Hprt) using the ΔΔCT method. Primers used see SI table 5

#### Dorsal root ganglion culture

DRG neurons were quickly and carefully dissected from the vertebral column^11^ and enzymatically digested for 75-90mins at 37°C 5% CO_2_ (collagenase type II - 4mg/ml, dispase type II – 4.7 mg/ml in Hank’s balanced salt solution Mg^2+^ and Ca^2+^ free (HBSS-Gibco^TM^ ThermoFisher Scientific)). Cells were then mechanically dissociated using fire polished glass pipettes plated into wells of a 24-well plate containing Poly-D-lysine/Laminin coated glass coverslips. Cells were maintained in supplemented Neurobasal^TM^ medium (Gibco^TM^ ThemroFisher Scientific) (2% (v/v) B27, 1%(v/v) GlutaMax, 1%(v/v) Antibiotic-antimycotic, Mouse nerve growth factor (50 ng/ml; NGF, PeproTech) and 10 ng/ml glial-derived neurotrophic factor (GDNF, PeproTech). Cells were used/analysed 24-72 hrs later.

### DRG electroporation

Electroporation was performed on dissociated cells prior to plating. Neurons were resuspended in 10 ul Buffer R containing 1 ug of plasmid DNA. Neurons were immediately withdrawn with an electroporation pipette/tip, an inserted into a Neon transfection system (ThermoFisher Scientific). Neurons were transfected using the following protocol, three 1500-V pulses of 10-ms duration. Cells were used/analysed 24-72 hrs later.

### Whole-cell patch-clamp

4 independent patch-clamp experiments were conducted using 4 (2 males 2 females) WT and 4 (2 males 2 females) *SLC45A4* KO mice. Prior to patch-clamp, an IB4-conjugated Alexa 488 live dye was added to the cells for 30 mins at 37°C to distinguish IB4-binding/positive and IB4-non binding/negative neurons. Using this we defined small (diameter < 25 µm) IB4-positive neurons as non-peptidergic nociceptors and small IB4-negative neurons as peptidergic nociceptors. Axopatch 200B amplifier and Digidata 1550 acquisition system (Molecular Devices, UK) were used an experiments performed at room temperature. Data were sampled at 20 kHz and low-pass filtered at 5 kHz. Series resistance was compensated 80 to 85% to reduce voltage errors. Alexa 488+ neurons were detected with an Olympus microscope with an inbuilt GFP (green fluorescent protein) filter set (470/40x excitation filter, dichroic LP 495 mirror, and 525/50 emission filter). Filamental borosilicate glass capillaries (1.5 mm OD, 0.84 mm ID; World Precision Instruments, United Kingdom) were pulled to form patch pipettes of 2 to 5 MΩ tip resistance and filled with an internal solution containing (mM): 100 K+ gluconate, 28 KCl, 1 MgCl_2_, 5 MgATP, 10 HEPES, and 0.5 EGTA; pH was adjusted to 7.3 with KOH and osmolarity set at 305 mOsm (using glucose). Neurons were maintained in a chamber constantly perfused with a standard extracellular solution containing (mM): 140 NaCl, 4.7 KCl, 2.5 CaCl_2_, 1.2 MgCl_2_, 10 HEPES, and 10 glucose; pH was adjusted to 7.3 with NaOH and osmolarity set at 315 mOsm (using glucose). There was a calculated −13 mV junction potential when using these solutions; voltage values were adjusted to compensate for this.

Resting membrane potential (RMP) was measured in bridge mode (I = 0). In current-clamp mode, neurons were held at −60 mV. Input resistance was derived by measuring the change in membrane voltage caused by a 80 pA hyperpolarising current step. Rheobase was determined by applying 50 ms depolarising current steps, (Δ10 pA), until action potential (AP) generation. Suprathreshold activity/repetitive firing was assed using two protocols. First, 500 ms depolarising current steps (Δ50 pA) until a final step current of 950 pA. Second, using a ramp protocol that gradually injects current from 0-1nA, with the duration of the ramp increasing (Δ100 ms) each time until a final ramp stimulus of 1000 ms. All data were analysed by Clampfit 10 software (Molecular Devices).

### Animal behaviour

A total of 15 (8 male, 7 female) wild type mice, 14 (7 male, 7 female) heterozygous mice, and 7 (3 male, 4 female) homozygous knockout mice were used for behavioural studies. All mice were littermates, age, sex matched where possible. All mice included in the behavioural pipelines underwent all behavioural tests. Mice were tested at consistent time of day during the light phase, in the same environment by the same experimenter. Mice were habituated to their testing environments and equipment before testing days. The experimenter was blind to animal genotype until post behavioural analysis was complete. Mice were randomly assigned a test environment and test order, which was achieved by random selection from their home cages. Unless otherwise stated, each test was conducted three times on different days to obtain an average baseline score.

#### Open field

Each mouse was individually placed into an open black testing box (size) with a grid marked into the floor. Mice were allowed to freely explore the open field for 3 min while being video recorded and tracked using ANY maze software. The following parameters were measured; number of rears, number of gridline crossing, total distance travelled and max speed when travelling.

#### Rotarod

Mice were placed onto a speed controlled horizontal rod/beam (Ugo Basile). Once mice were placed onto the stationary rod, an increasing speed (ramp) protocol was applied (0.5 RPM/s for 20 s, 0.25 RPM/s for 160 s, 0.16 RPM/s for 120 s to reach a final speed of 80 RPM/s). Mice were monitored until they fell from the rotating rod. Latency to fall (s) and final speed (rpm) was recorded, on two testing days and averaged.

#### Mechanical testing

Mice were placed into a testing box (5 x 5 x 10 cm) elevated on a wire mesh base and mice allowed to acclimatise for 30-60 min. The plantar hind paws were tested using von Frey hairs (up-down method^12^) to calculate the 50% paw withdrawal threshold, or a pin prick attached to a 1 g von Frey hair to measure the latency to withdraw (using a high frame rate camera).

#### Thermal sensory testing

Mice were plated onto a Perspex enclosed hot plate that was either set to 48°C, 50°C or 53°C. Mice were observed and timed until their hind paws reacted to the hot plate (Lifting, flicking, licking, cut off set to 20 s). The latency to respond to each hot plate was measured. For the Hargreaves test mice were placed into a test box (5 x 5 x 10 cm) on elevated glass and acclimatised for 30-60 min. The Hargreaves radiant heat source was applied to each paw three times and latency to withdraw was recorded. Noxious cold sensitivity was measured using the dry ice test. Mice were placed in test boxes, and elevated on a 5mm thick glass platform. Dry ice pieces were placed into a 2 ml syringe (top removed), which was placed against the glass under a visible hind paw. Latency to withdraw from the dry ice/glass was measured and three measurements were taken for each hind paw.

### Thermal gradient test

Mice were allowed to freely explore a thermal gradient apparatus (BIOSEB) which consisted of a metal platform that was heated at one end and cooled at the other. This created a thermal gradient platform the ranged from 54°C – 6°C. During the 60 min exploration mice were video tracked using a HD webcam and ANYMaze software, and time in different temperature zones analysed. This test was only performed once to avoid learning confounds. One heterozygous mouse (female) was excluded from this test due to a camera fault during the 60 min run.

### Tonic pain assay – Formalin

Mice received a single subcutaneous injection of 20 ul of 2% formalin in the left hind paw. Mice were immediately placed into a test box (5 x 5 x 10 cm), elevated on a glass platform, surrounded by angled mirrors, all above a video camera. Mice were video recorded for 1 hr while the experimenter left the room. The videos were analysed offline by two blinded experimenters, each measured the duration of nocifensive behaviours (lifting, licking, flinching, shaking) of the injected hind paw every 5 min for 1 hr. The first formalin phase was defined as 0-15 min, and the second phase was defined as 15-60 min.

### Statistical analysis and reproducibility

#### Animal work

Data were analysed using GraphPad Prism 10 and ImageJ/Fiji (NIH). The number of samples and statistical test used for each experiment have been included in the figure legends. In the case where n equals animal, the number of cells have also been described. In all histology experiments, at least 3 sections containing cells were analysed per mouse. The number of animals used in behavioural experiments have been described in the legends and the relevant methods section.

Statistical analysis used were unpaired t-test, one-way ANOVA with Tukey multiple comparisons test, extra sum of squares F-test, Kruskal-Wallis test with Dunn’s multiple comparisons test, two-way ANOVA with Holm-Sidak’s multiple comparisons test, repeated measures two-way ANOVA with Holm-Sidak’s multiple comparisons test. Exact n numbers and P values have been reported in figure legends. Full statistics for each test used in the study can be found here. **Fig. 3f** Latency to fall one way-ANOVA, F = 4.581, P = 0.0176, DFn, DFd = 2, 33. WT vs KO, P = 0.0132, q = 4.267, DF. = 33. Final speed one-way ANOVA, F = 5.094, P = 0.0118, DFn, DFd. = 2, 33. WT vs KO, P = 0.0085, q = 4.513. DF = 33. **Fig. 3g** One-way ANOVA, F = 0.1043, P = 0.9013, DFn, DFd. = 2, 33. **Fig. 3h** one-way ANOVA, F = 7.824. P = 0.0017, DFn, DFd. = 2, 33. WT vs KO, P = 0.002, q = 5.271, DF = 33. HET vs KO, P = 0.0039, q = 4.938, DF = 33. **Fig. 3i** one-way ANOVA, F = 3.703, P = 0.0354, DFn, DFd. = 2, 33. WT vs KO, P = 0.0398, q = 3.615, DF = 33. HET vs KO, P = 0.0546, q = 3.413, DF = 33. **Fig. 3j** two-way ANOVA, Holm-Sidak’s test, WT vs HET at 29°C, P = 0.0096, t = 2.959, DF = 544. WT vs KO at 29°C, P = 0.0275, t = 2.469, DF = 544. **Fig. 3k** Extra sum of squares F test, F = 2.44, P = 0.0242, DFn, DFd = 6, 586. **Fig. 3l** RM two-way ANOVA, Holm-Sidak’s test, WT vs KO at 5 mins, P = 0.0113, t = 3.298, DF = 19.05. HET vs KO at 5 mins, P = 0.0276, t = 2.734, DF = 17.41. HET vs KO at 10 mins, P = 0.0182, t = 3.237, DF 13.75. **Fig. 3m** Phase 1: Kruskal-Wallis test, P = 0.0208, KW = 7.743. Dunn’s WT vs KO, P = 0.020, Z = 2.713. Phase 2: ANOVA, F = 0.4275, P = 0.6557, DFn, DFd = 2, 33. **Fig 4b**, two-way ANOVA, Holm-Sidak test, WT vs KO, CGRP: P = 0.4809, t = 1.492, DF = 20. IB4: P = 0.7123, t = 1.140, DF = 20. NF200: P = 0.6926, t = 1.171, DF = 20. TH: P = 0.8667, t = 0.0678, DF 20. **Fig 4d**, unpaired t-test, P = 0.758, t = 0.329, DF = 4. **Fig 4f**, Kruskal-Wallis test, P = 0.559, KW = 2.065. Dunn’s IB4-postive WT vs KO, P = >0.99, Z = 0.437. IB4-negative WT vs KO, P = 0.677, Z = 0.957. **Fig 4g**, RM two-way ANOVA, Holm-Sidak’s test, WT vs KO 50-950 pA, P > 0.95, t = 0.007-1.44, DF = 323. **Fig 4h**, RM two-way ANOVA, Holm-Sidak’s test, WT vs KO 100-1000 ms, P > 0.99, t = 0.011-0.655, DF = 150. **Fig 4i**, RM two-way ANOVA, Holm-Sidak’s test, WT vs KO at 100 pA, P = 0.039, t 3.038, DF = 418, at 150 pA, P = 0.0027, t = 3.807, DF = 418, at 200 pA, P = 0.0002, t = 4.507, DF = 418, at 250 pA, P = 0.0006, t = 4.212, DF = 418, at 300 pA, P = 0.047, t = 2.961, DF = 418. **Fig 4j**, RM two-way ANOVA, Holm-Sidak’s test, WT vs KO at 300 ms, P = 0.036, t = 2.623, DF = 220, at 400 ms, P = 0.021, t = 2.904, DF = 220, at 500 ms, P = 0.011, t = 3.314, DF = 220, at 600 ms, P = 0.013, t = 3.22, DF = 220, at 700 ms, P = 0.021, t = 2.998, DF = 220, at 800 ms, P = 0.015, t = 3.127, DF = 220, at 900 ms, P = 0.021, t = 2.951, DF = 220, at 1000 ms, P = 0.037, t = 2.623, DF = 220. **Extended data figure 7d,** DRG: one-way ANOVA, F = 305.7, P < 0.0001, DFn, DFd = 2, 10. WT vs HET, P < 0.0001, q = 21.28, DF = 10. WT vs KO, P < 0.0001, q = 34.32, DF = 10. HET vs KO, P < 0.0001, q = 12.37, DF = 10. Spinal cord: one-way ANOVA, F = 34.51, P < 0.0001, DFn, DFd = 2, 9. WT vs HET, P = 0.0262, q = 4.535, DF = 9. WT vs KO, P < 0.0001, q = 11.65, DF = 9. HET vs KO, P = 0.0018, q = 7.118, DF = 9. Brain: one-way ANOVA, F = 16.23, P = 0.001, DFn, DFd = 2, 9. WT vs HET, P = 0.1206, q = 3.142, DF = 9. WT vs KO, P = 0.0008, q = 7.996, DF = 9. HET vs KO, P = 0.0185, q = 4.854, DF = 9. **Extended data figure 7e,** one-way ANOVA, F = 980.5, P < 0.0001, DFn, DFd = 2, 3749. WT vs HET, P < 0.0001, q = 38.58, DF = 3749. WT vs KO, P < 0.0001, q = 61.59, DF = 3749. HET vs KO, P < 0.0001, q = 27.54, DF = 3749. **Extended data figure 8b,** one-way ANOVA, F = 0.2183, P = 0.8050, DFn, Dfd = 2, 33. **Extended da**ta figure 8c, one-way ANOVA, F = 2.308, P = 0.1163, DFn, Dfd = 2, 33. **Extended data figure 8d** one-way ANOVA, F = 0.6064, P = 0.5513, DFn, Dfd = 2, 33. **Extended** data figure 8e one-way ANOVA, F = 0.3041, P = 0.7398, DFn, Dfd = 2, 33. **Extended data figure 8f** one-way ANOVA, F = 2.012, P = 0.1497, DFn, Dfd = 2, 33. **Extended data figure 8g** one-way ANOVA, F = 1.035, P = 0.3666, DFn, Dfd = 2, 33. **Extended data figure 8h** one-way ANOVA, F = 2.960, P = 0.0657, DFn, Dfd = 2, 33. **Extended data figure 8i** one-way ANOVA, F = 0.7461, P = 0.4820, DFn, Dfd = 2, 33. **Extended data figure 9b,** no. fragments, one-way ANOVA, F = 0.700, P = 0.5283, DFn, Dfd = 2, 7. NMJ area, one-way ANOVA, F = 0.4309, P = 0.666, DFn, Dfd = 2, 7. α-BuTx/synapNF, one-way ANOVA, F = 1.197, P = 0.3572, DFn, Dfd = 2, 7. synapNF /α-BuTx, one-way ANOVA, F = 1.587, P = 0.2703, DFn, Dfd = 2, 7.

## Data availability

Genotype data, imputed data and linked phenotypes for the UK Biobank dataset, can be accessed via the UKB research analysis platform (RAP) (https://ukbiobank.dnanexus.com/landing). The UK Biobank Resource was used under application number 49572. Structural data has been deposited in wwPDB, EMD-51377, 9GIU, and EMD-51365, 9GHZ. Datasets relating to studies in mouse may be obtained from the corresponding author.

## Acknowledgements

This work was funded by: Wellcome Trust (Wellcome Investigator Grants to DB, 223149/Z/21/Z and SN 215519/Z/19/Z & 219531/Z/19/Z); SM is funded through a Wellcome graduate studentship; UK Medical Research Council (grant ref. MR/T020113/1 to DB SJM); Diabetes UK (grant agreement19/0005984 to GB DLB); MRC and Versus Arthritis (to the PAINSTORM consortium as part of the Advanced Pain Discovery Platform MR/W002388/1); European Union Horizon 2020 research and innovation programme (grant agreement No 633491 to DOLORisk); National Institute for Health and Care Research (NIHR) Oxford Health Biomedical Research Centre. This research was supported in part by the Intramural Research Program of the NIH (SML), and by US Department of Veterans Affairs (Merit Award I01 BX004820) and NIH grant R01 AA030056. The views expressed are those of the author(s) and not necessarily those of the NIHR, NIH or the Department of Health and Social Care. For the purpose of open access, the author has applied a CC BY public copyright license to any Author Accepted Manuscript version arising from this submission.

## Author contributions

Conceptualization: BHS, DLB, GB, HH, JCD, JEL, JLP, JPS, MA, MT, SJM, SM, SML, SN, VL, WL, YYD

Methodology: AF, BHS, GB, GK, HLH, HRK, JCD, JPS, MA, MT, PS, SJM, SM, SM, SML, ST, SZ, WL

Investigation: AF, GB, GK, JCD, JLP, JPS, MA, MT, PS, SJM, SM, SM, ST, SZ, WL

Visualization: DLB, GK, JCD, JLP, MA, MT, PS, SJM, SM, SM, SN, SZ, VL, WL, YYD

Funding acquisition: BHS, DLB, HRK, JLP, SJM, SM, SML, SN, VL

Project administration: BHS, DLB, GB, JLP, MA, SJM, SM, SN, VL, WL

Writing – original draft: DLB, GB, JLP, MA, SJM, SM, SN, VL, WL

Review & editing: All authors

## Materials & Correspondence

Correspondence should directed to either:

David L Bennett, The Nuffield Department of Clinical Neuroscience, University of Oxford, The John Radcliffe Hospital, Headley Way Oxford, UK OX39DU,

Simon Newstead, Department of Biochemistry, University of Oxford, Oxford OX1 3QU, UK. The Kavli Institute for Nanoscience Discovery, University of Oxford, Oxford, OX1 3QU, UK

## Supplementary information

This manuscript includes supplementary methods, supplementary tables 1-6, and supplementary files.

