## Supplementary Material for "Identification of SLC45A4 as a pain gene encoding a neuronal polyamine transporter"

#### Extended data figures

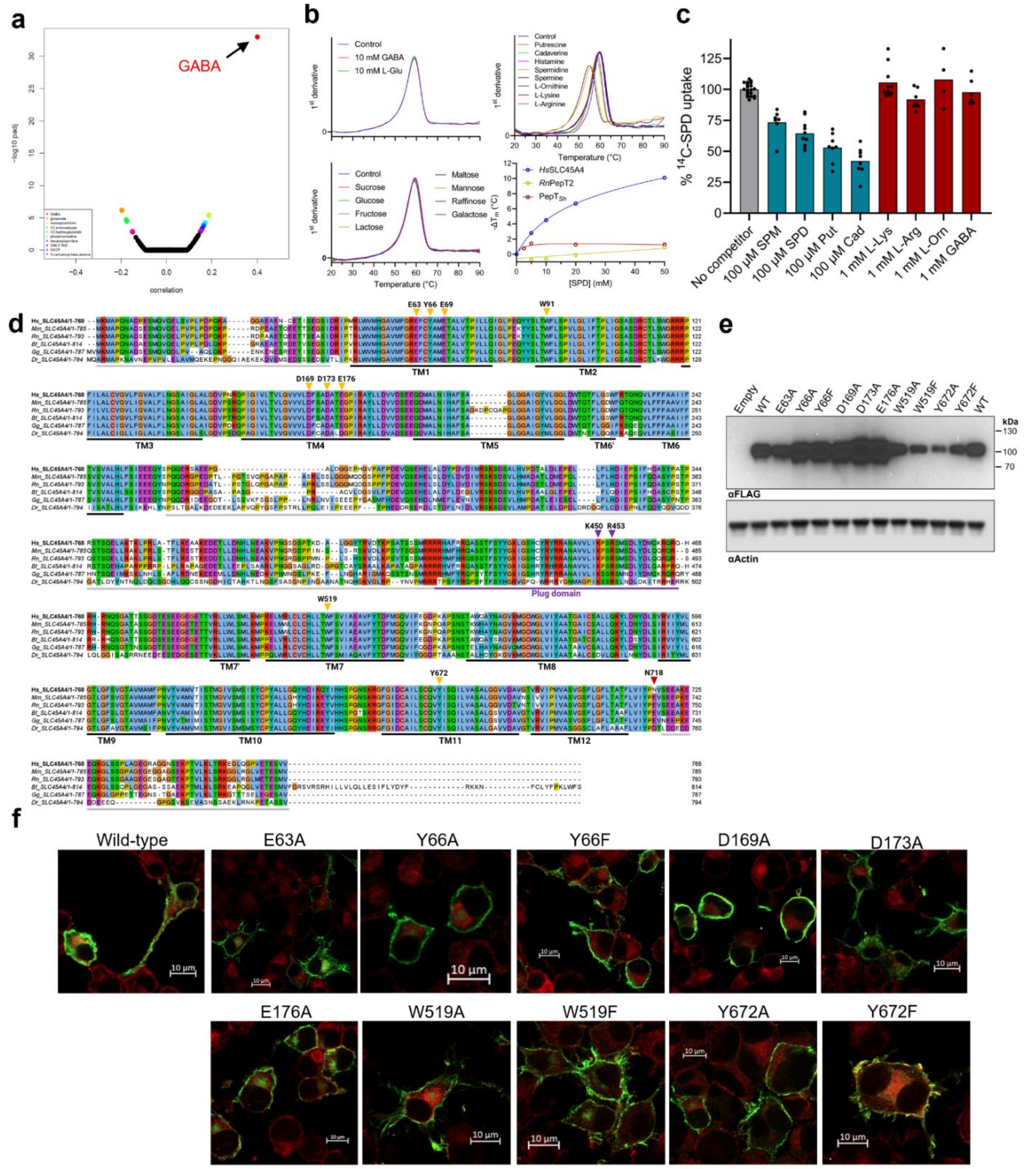

**Extended Data Fig. 1. Functional analysis of human SLC45A4. a,** Correlation analysis between metabolomics and expression datasets identified GABA as being positively associated with overexpression of SLC45A4. **b,** Effect of different metabolites on the thermal stability of SLC45A4 in detergent. **c,** Competition of $^{14}C$-SPD transport by WT *Hs*SLC45A4 in Neuro-2A cells showed inhibition with polyamines, but no recognition of cationic amino acids or GABA. **d,** Sequence alignment of human (*Hs*; Q5BKX6), mouse (*Mm*, Q0P5V9), rat (*Rn*, D4ADC6), bovine (*Bt*, E1BGZ7), chicken (*Gg*, A0A1D5NU91) and zebrafish (*Dr*, E9QH03) SLC45A4 sequences. Secondary structure, as observed in the cryo-EM structure of SLC45A4 and residues shown to be important to function are highlighted. Regions not resolved in the cryo-EM maps are highlighted in grey. **e,** Western blots confirming expression of mutants in N2A cells. **f,** Confocal microscopy shows localisation of SLC45A4 (Green) in the plasma membrane as determined by co-localisation with the $Na^{+}/K^{+}$ ATPase (red).

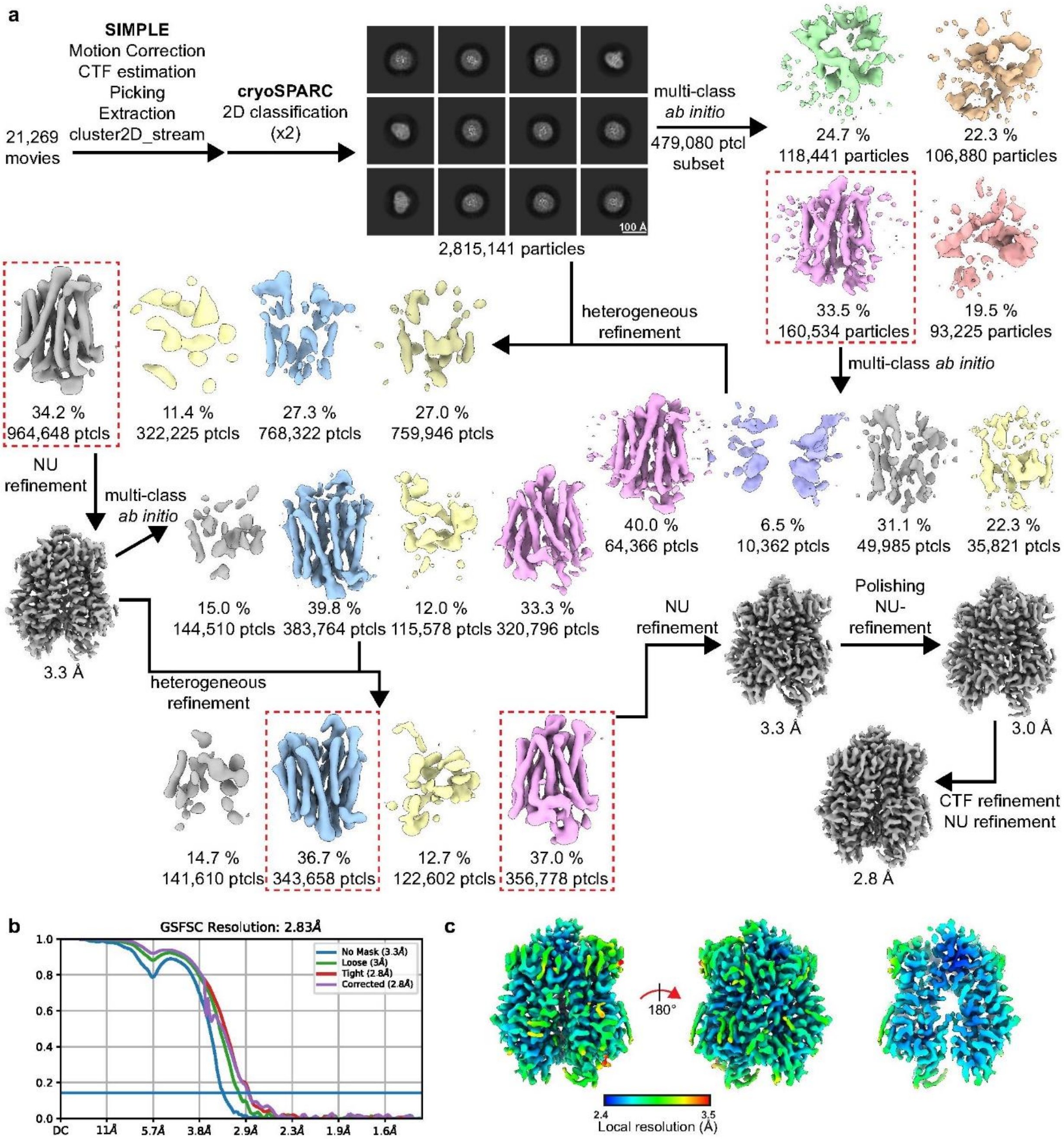
a
SIMPLE
Motion Correction
CTF estimation
Picking
Extraction
cluster2D_stream
21,269 movies
cryoSPARC
2D classification (x2)
100 Å
2,815,141 particles
multi-class ab initio
479,080 ptcl subset
24.7 %
118,441 particles
22.3 %
106,880 particles
33.5 %
160,534 particles
19.5 %
93,225 particles
multi-class ab initio
heterogeneous refinement
34.2 %
964,648 ptcls
11.4 %
322,225 ptcls
27.3 %
768,322 ptcls
27.0 %
759,946 ptcls
40.0 %
64,366 ptcls
6.5 %
10,362 ptcls
31.1 %
49,985 ptcls
22.3 %
35,821 ptcls
NU refinement
multi-class ab initio
3.3 Å
15.0 %
144,510 ptcls
39.8 %
383,764 ptcls
12.0 %
115,578 ptcls
33.3 %
320,796 ptcls
heterogeneous refinement
NU refinement
Polishing
NU-refinement
3.3 Å
3.0 Å
14.7 %
141,610 ptcls
36.7 %
343,658 ptcls
12.7 %
122,602 ptcls
37.0 %
356,778 ptcls
CTF refinement
NU refinement
2.8 Å
b
GSFSC Resolution: 2.83Å
No Mask (3.3Å)
Loose (3Å)
Tight (2.8Å)
Corrected (2.8Å)
1.0
0.8
0.6
0.4
0.2
0.0
DC
11Å
5.7Å
3.8Å
2.9Å
2.3Å
1.9Å
1.6Å
c
180°
2.4
3.5
Local resolution (Å)

**Extended Data Fig. 2. Cryo-EM processing workflow of SLC45A4 in LMNG, including local and global resolution estimates**. **a,** Image processing workflow. **b,** Gold-standard Fourier Shell Correlation (FSC) curves for global resolution estimation. **c,** Local resolution estimate of the volume.

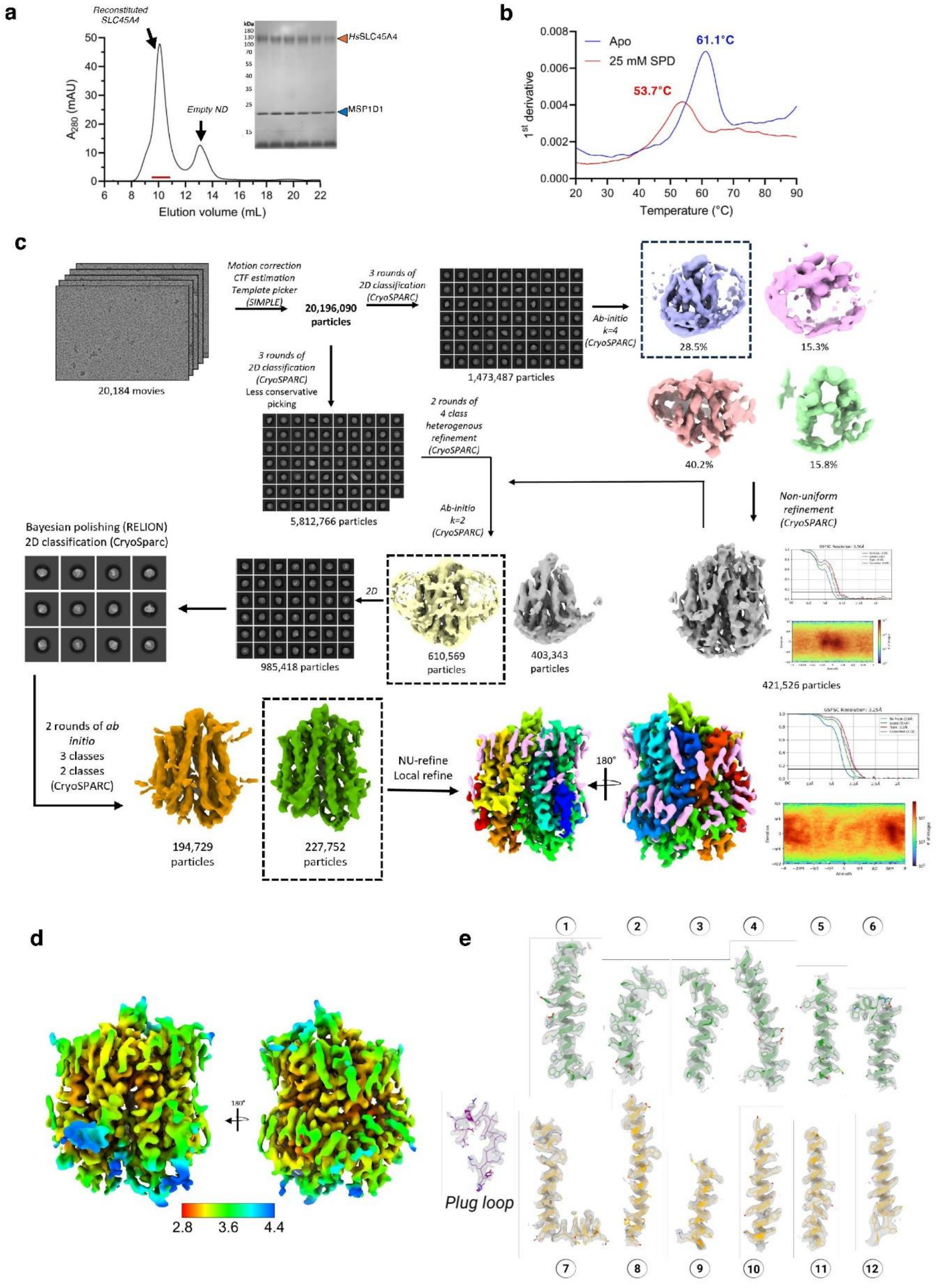
a
Reconstituted SLC45A4
Empty ND
HsSLC45A4
MSP1D1
A280 (mAU)
Elution volume (mL)
b
Apo
25 mM SPD
61.1°C
53.7°C
1st derivative
Temperature (°C)
c
Motion correction
CTF estimation
Template picker
(SIMPLE)
20,196,090 particles
3 rounds of 2D classification (CryoSPARC)
1,473,487 particles
Ab-initio k=4 (CryoSPARC)
28.5%
15.3%
40.2%
15.8%
20,184 movies
3 rounds of 2D classification (CryoSPARC) Less conservative picking
2 rounds of 4 class heterogenous refinement (CryoSPARC)
5,812,766 particles
Ab-initio k=2 (CryoSPARC)
Non-uniform refinement (CryoSPARC)
Bayesian polishing (RELION)
2D classification (CryoSparc)
2D
985,418 particles
610,569 particles
403,343 particles
421,526 particles
2 rounds of ab initio
3 classes
2 classes
(CryoSPARC)
194,729 particles
227,752 particles
NU-refine
Local refine
180°
d
2.8
3.6
4.4
e
Plug loop
1
2
3
4
5
6
7
8
9
10
11
12

**Extended Data Fig. 3. Nanodisc reconstitution, CryoEM data acquisition and processing**. **a** Reconstitution of HsSLC45A4, purified from HEK293F, into EBC:MSP1D1 nanodiscs. **b** NanoDSF scan (1st derivative) of SLC45A4 nanodiscs with and without 25 mM SPD, showing similar destabilisation as observed in detergent. **c** Overview of data acquisition and processing of the SLC45A4 nanodisc sample, resulting in a well-resolved map with a global resolution of 3.25 Å. Density corresponding to protein is shown in rainbow, and lipid/detergent in pink. **d** Local resolution map. **e** Electron density of the transmembrane helices and plug domain loop.

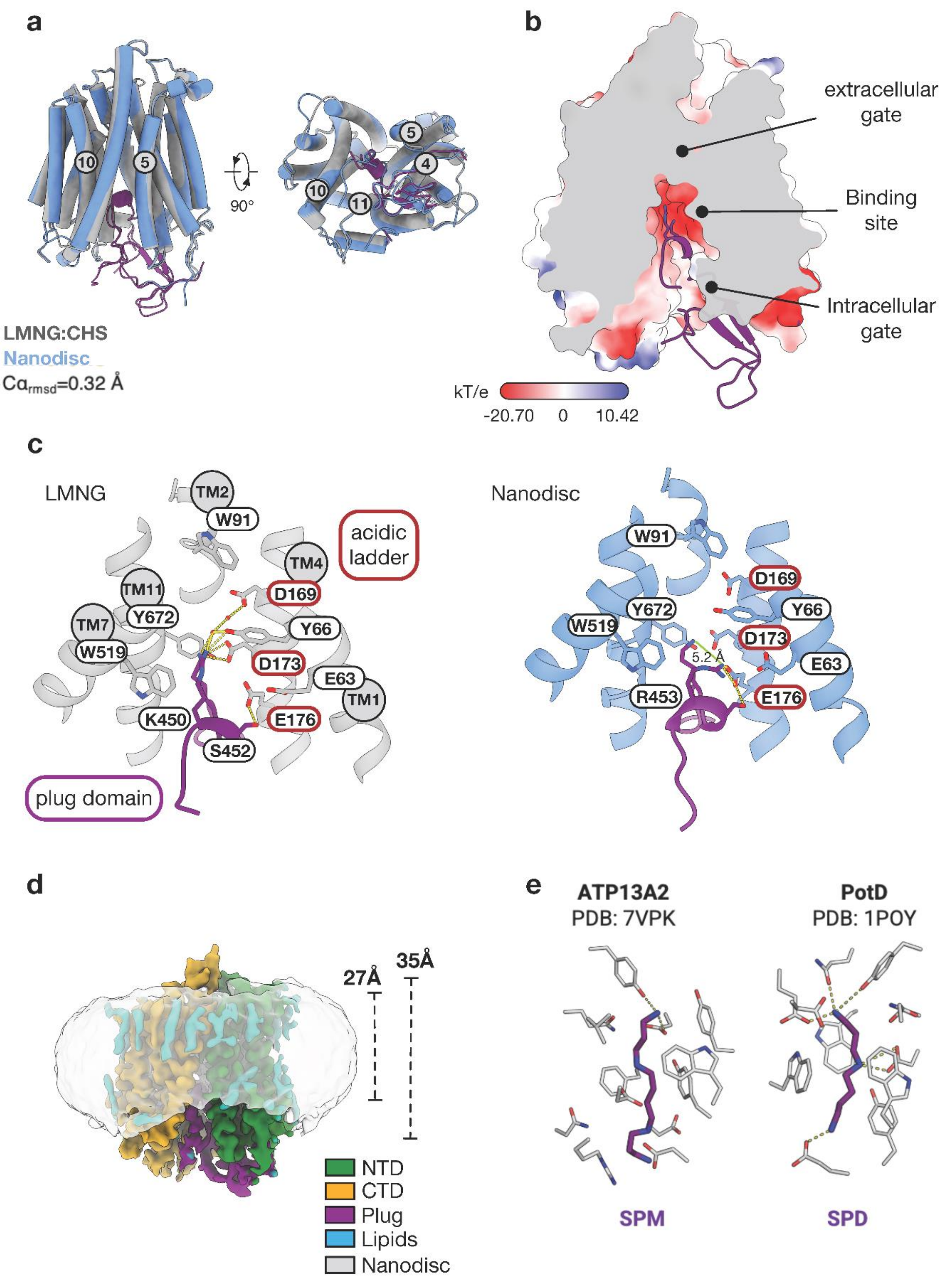
a
10
5
90°
10
11
4
5
LMNG:CHS
Nanodisc
Cα$_{rmsd}$=0.32 Å
b
extracellular gate
Binding site
Intracellular gate
kT/e
-20.70
0
10.42
c
LMNG
TM2
W91
acidic ladder
TM4
TM11
D169
TM7
Y672
Y66
W519
D173
E63
TM1
K450
E176
S452
plug domain
Nanodisc
W91
D169
Y672
Y66
W519
D173
5.2 Å
E63
R453
E176
d
27Å
35Å
NTD
CTD
Plug
Lipids
Nanodisc
e
ATP13A2
PDB: 7VPK
PotD
PDB: 1POY
SPM
SPD

**Extended Data Fig. 4. Structural analysis of SLC45A4. a,** Superposition of the SLC45A4 structures determined in LMNG and nanodisc. Gating helices are labelled. **b,** Slice through the electrostatic surface of SLC45A4 showing the location of the plug domain relative to the canonical MFS binding site. The sealed extracellular gate is indicated. **c,** View of the canonical MFS binding site showing the main polar interactions formed between Lys450, Ser452 and Arg453 on the plug domain with the side chains on the transmembrane helices. We have used these interactions as a proxy for polyamine recognition. Left hand panel shows the structure from the LMNG sample (grey helices), the right-hand panel the nanodisc structure (blue helices) with altered rotamer position for Arg453. **d,** Cryo-EM density of the SLC45A4 nanodisc structure contoured at a threshold level of 0.2 for the protein molecule (B-factor sharpened map) and 0.07 for the nanodisc (unsharpened map). **e,** Binding site for SPD in ATP13A2 and PoTD respectively, showing the main polar interactions (dashed lines)**.**

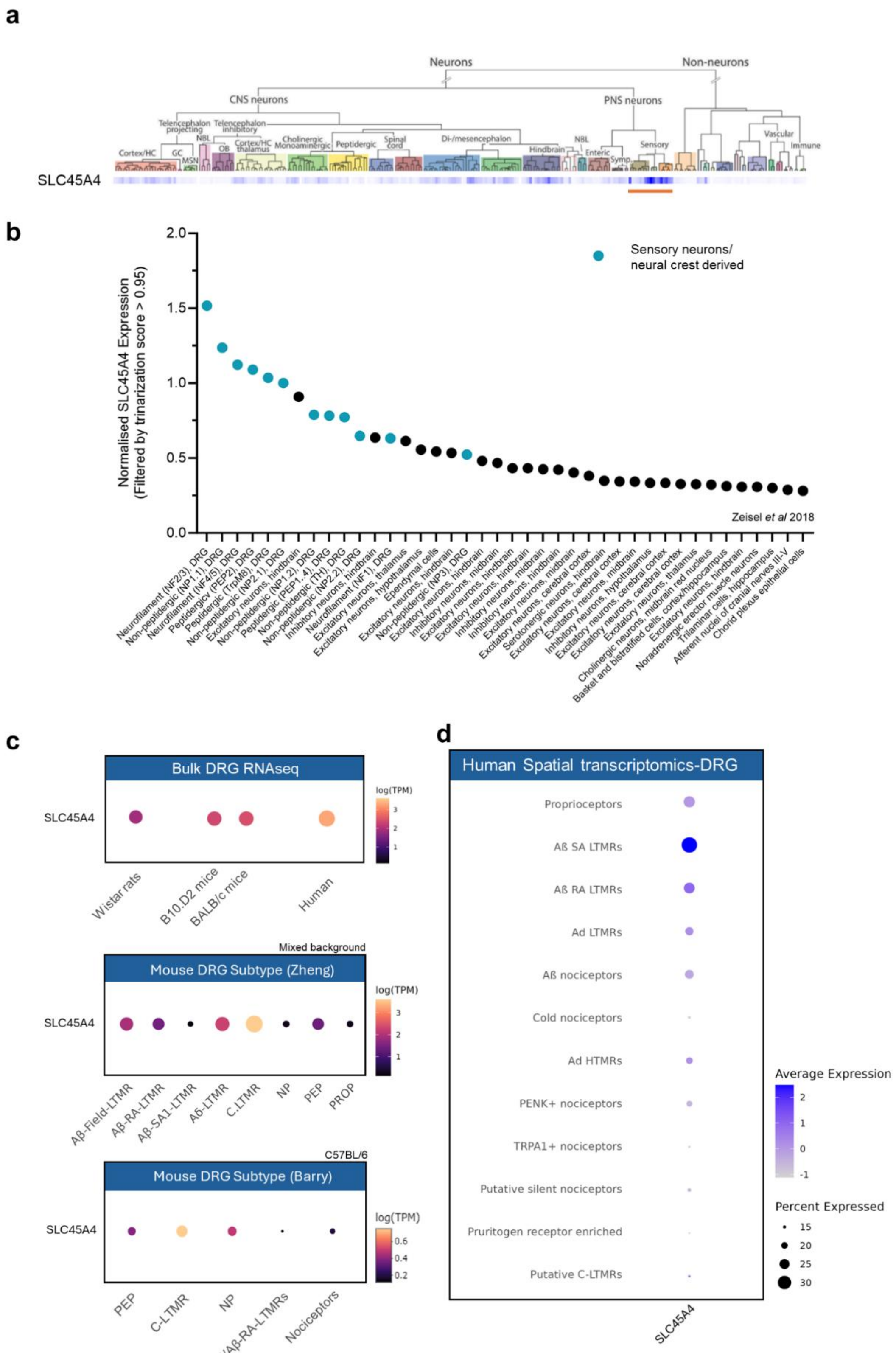


1324

1325

1326

1327

**Extended data Fig. 5. SLC45A4 expression in mouse and human sensory neurons evidenced in multiple data sets. a,** Heat map of *SLC45A4* mRNA expression detected using single cell sequencing of the entire mouse nervous system (high expression in blue, no expression in white). Compared across the nervous system, *SLC45A4* mRNA is enriched in sensory neuron subpopulations (red bar). Data from Mousebrain.org,[35]. **b,** Filtering the same data set based on a trimerization score >0.95 (increased confidence of expression), 40 regions were identified, 12 regions were identified as sensory neuron subtypes and neural crest derived. 9 out of the top 10 ten regions were sensory neuron subtypes of the DRG[35]. **c,** Top: Bulk DRG RNA sequencing of different rodent species, strains and human DRG. *SLC45A4* mRNA was detected at high levels in across rats, mouse and human DRG. Middle: Deep sequencing of 8 mouse (mixed back ground) sensory neuron subtypes[36] and Bottom: Deep sequencing of 5 mouse (C57BL6) sensory neurons subtypes[37]. Both illustrate wide expression of SLC45A4 *mRNA* in mouse sensory neurons. **d,** Data from spatial transcriptomics of human dorsal root ganglia. *SLC45A4* mRNA is expressed broadly in most subtypes including nociceptors and mechanoreceptors[38]. c and d were generated from the DRG-directory (https://livedataoxford.shinyapps.io/drg-directory/).

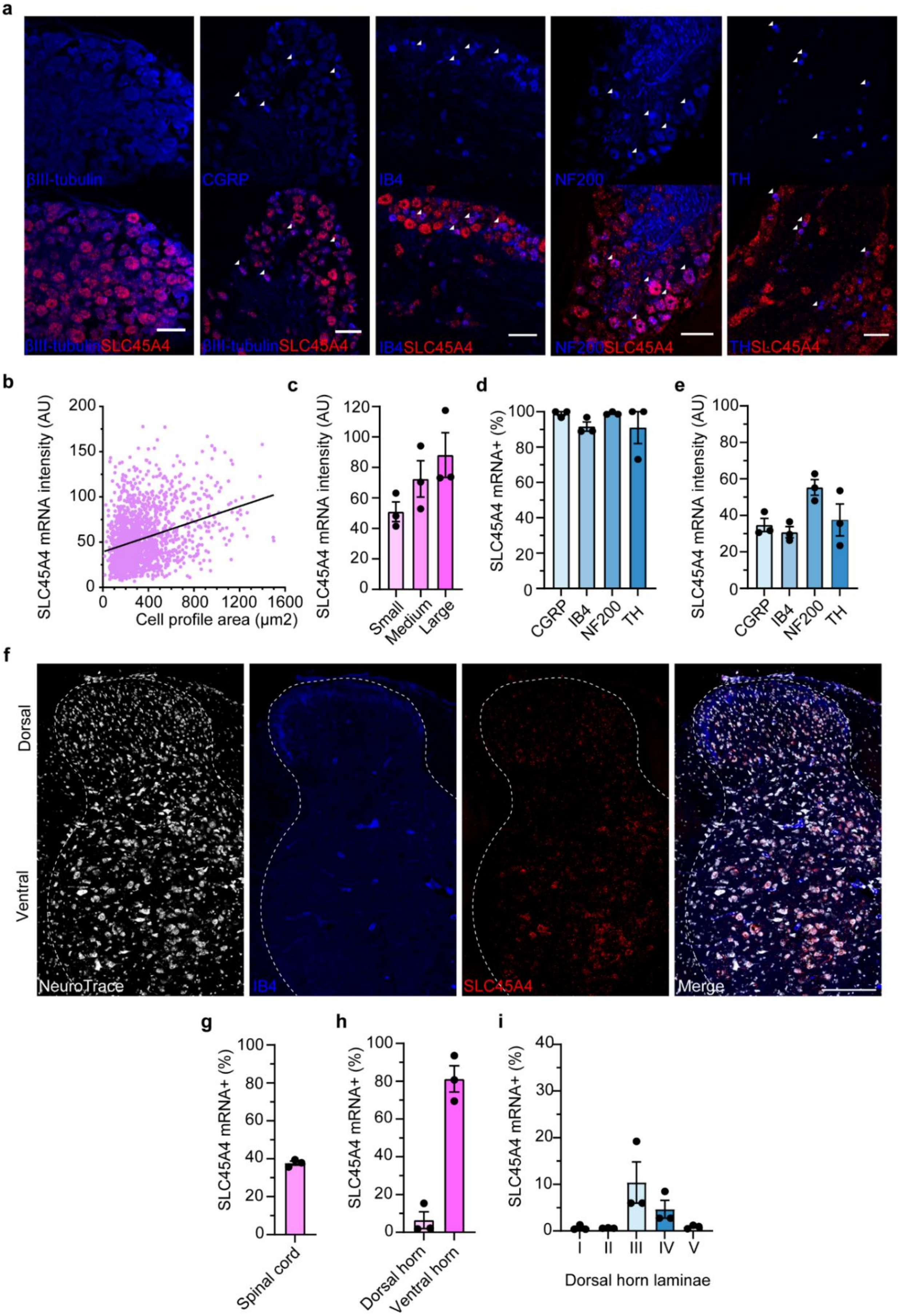
a
βIII-tubulin
CGRP
IB4
NF200
TH
βIII-tubulinSLC45A4
βIII-tubulinSLC45A4
IB4SLC45A4
NF200SLC45A4
THSLC45A4
b
SLC45A4 mRNA intensity (AU)
Cell profile area (μm2)
c
SLC45A4 mRNA intensity (AU)
Small
Medium
Large
d
SLC45A4 mRNA+ (%)
CGRP
IB4
NF200
TH
e
SLC45A4 mRNA intensity (AU)
CGRP
IB4
NF200
TH
f
Dorsal
Ventral
NeuroTrace
IB4
SLC45A4
Merge
g
SLC45A4 mRNA+ (%)
Spinal cord
h
SLC45A4 mRNA+ (%)
Dorsal horn
Ventral horn
i
SLC45A4 mRNA+ (%)
I
II
III
IV
V
Dorsal horn laminae

1358

**Extended data Fig. 6. *SLC45A4* mRNA is broadly expressed in mouse sensory neurons. a,** Example images of RNA scope ISH of *SLC45A4* mRNA and co-localisation with all sensory neurons (βIII-tubulin) and sensory neuron subtype markers (CGRP, IB4, NF200, TH). **b,** *SLC45A4* mRNA intensity vs cell profile area, line of best fit showing a positive correlation. (1,777 cells, from 3 mice). **c,** This data was subdivided into small medium and large cells with *SLC45A4* mRNA intensity increasing with size (n = 3 mice, 1,461 small, 269 medium, 41 large cells). **d,** Percentage of sensory neuron subtypes that colocalise and express *SLC45A4* mRNA. **e,** *SLC45A4* mRNA intensity in each sensory neuron subpopulation with the highest signal intensity in the NF200 population, (d+e: n = 3 mice, no. cells: 144 CGRP, 273 IB4, 114 NF200, 157 TH). Scale bars 100 µm. **f,** Example images of RNA scope ISH of *SLC45A4* mRNA in the lumbar spinal cord, all neurons (NeuroTrace), primary afferent terminals (IB4), and SLC45A4 mRNA. Scale bar 200 µm. **g**, Percentage of all spinal cord neurons that express *SLC45A4* mRNA (n = 3 mice, 14,286 cells) **h**, Percentage of dorsal and ventral horn neurons that are *SLC45A4* mRNA+ (n =3 mice, dorsal 13,179 cells, ventral 5,402 cells). **i,** Percentage of neurons from different dorsal horn laminae that express *SLC45A4* mRNA (n =3 mice, I 749 cells, II 1,649 cells, III 1,557 cells, IV 1,384 cells, V 2,047 cells).

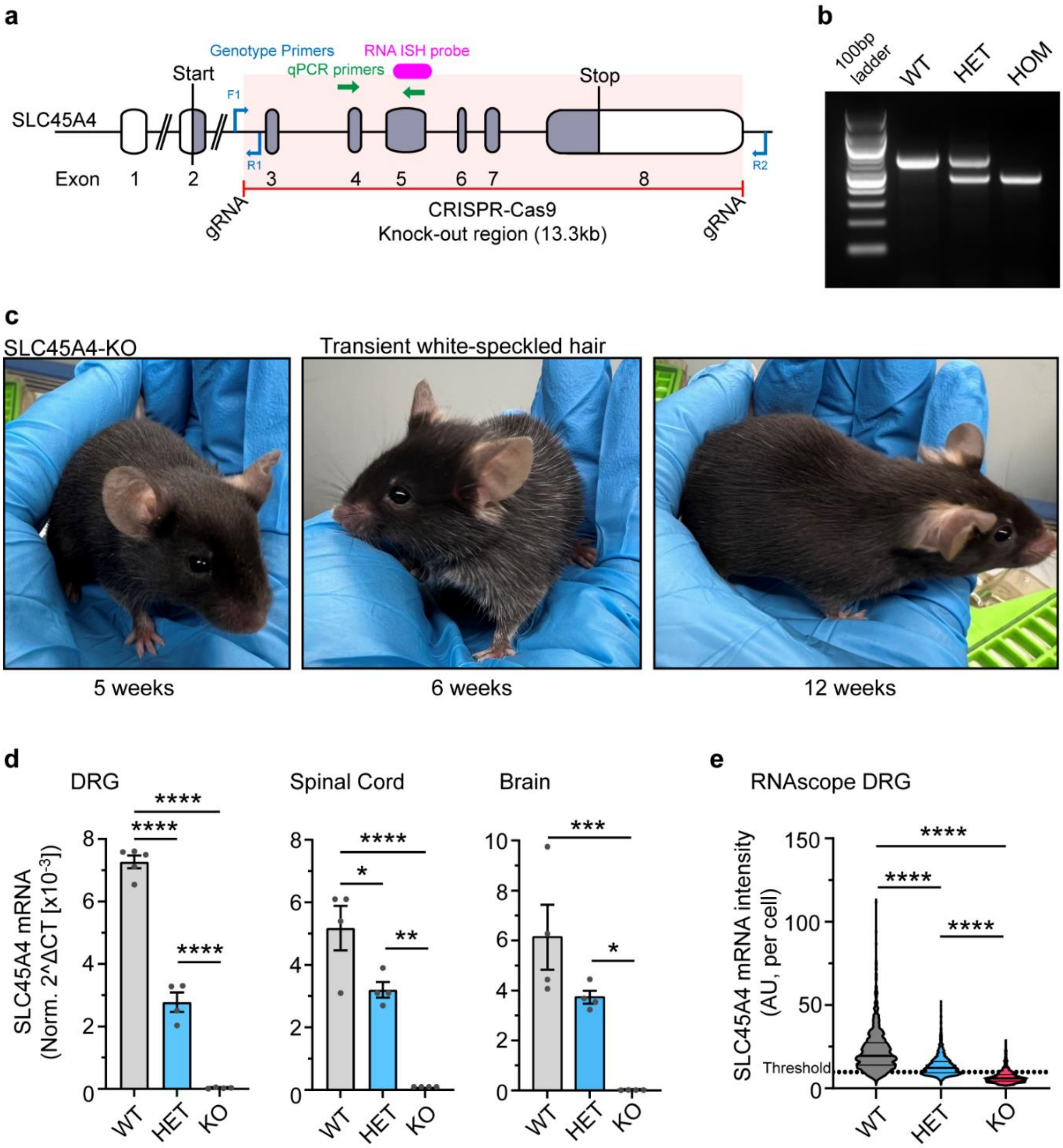
a
Genotype Primers
RNA ISH probe
qPCR primers
Start
Stop
F1
R1
R2
SLC45A4
Exon
1
2
3
4
5
6
7
8
gRNA
CRISPR-Cas9
Knock-out region (13.3kb)
gRNA
b
100bp ladder
WT
HET
HOM
c
SLC45A4-KO
Transient white-speckled hair
5 weeks
6 weeks
12 weeks
d
DRG
Spinal Cord
Brain
SLC45A4 mRNA (Norm. 2^ΔCT [x10-3])
WT
HET
KO
e
RNAscope DRG
SLC45A4 mRNA intensity (AU, per cell)
Threshold
WT
HET
KO

**Extended data Fig. 7. Generation and validation of an *SLC45A4* KO mouse. a,** Schematic of the *SLC45A4* locus outlining the 8 exons, the start and stop sites, the CRISPR-Cas9 KO region targeted with guide RNAs (gRNA). The illustration also highlights where the genotyping and qPCR primers target and the RNAscope probe. **b,** Example genotyping gel of WT, HET and KO mice. PCR amplified WT band 652bp, KO band 468bp. **c,** Images of *SLC45A4* KO mice at different ages, at about 6 weeks mice develop a salt and pepper/white speckled hair. This is transient and returns to normal by week 12. **d,** qPCR was used to analyse the expression of *SLC45A4* mRNA in WT HET and KO mice, in DRG, Spinal cord and Brain respectively. Heterozygous mice have significantly reduced *SLC45A4* mRNA compared to WT, and there is a complete absence of expression in KO mice across tissues (n = 4 mice per group) **e,** Analysis of *SLC45A4* mRNA signal intensity in DRG sections from WT, HET and KO mice (WT n = 1349 cells from 4 mice, HET n = 1452 cells from 3 mice, KO n = 951 cells from 3 mice). Threshold set using a negative control probe. d and e, one-way ANOVA with post hoc Tukey test, * $P < 0.05$, ** $P < 0.01$, *** $P < 0.001$, **** $P < 0.0001$.).

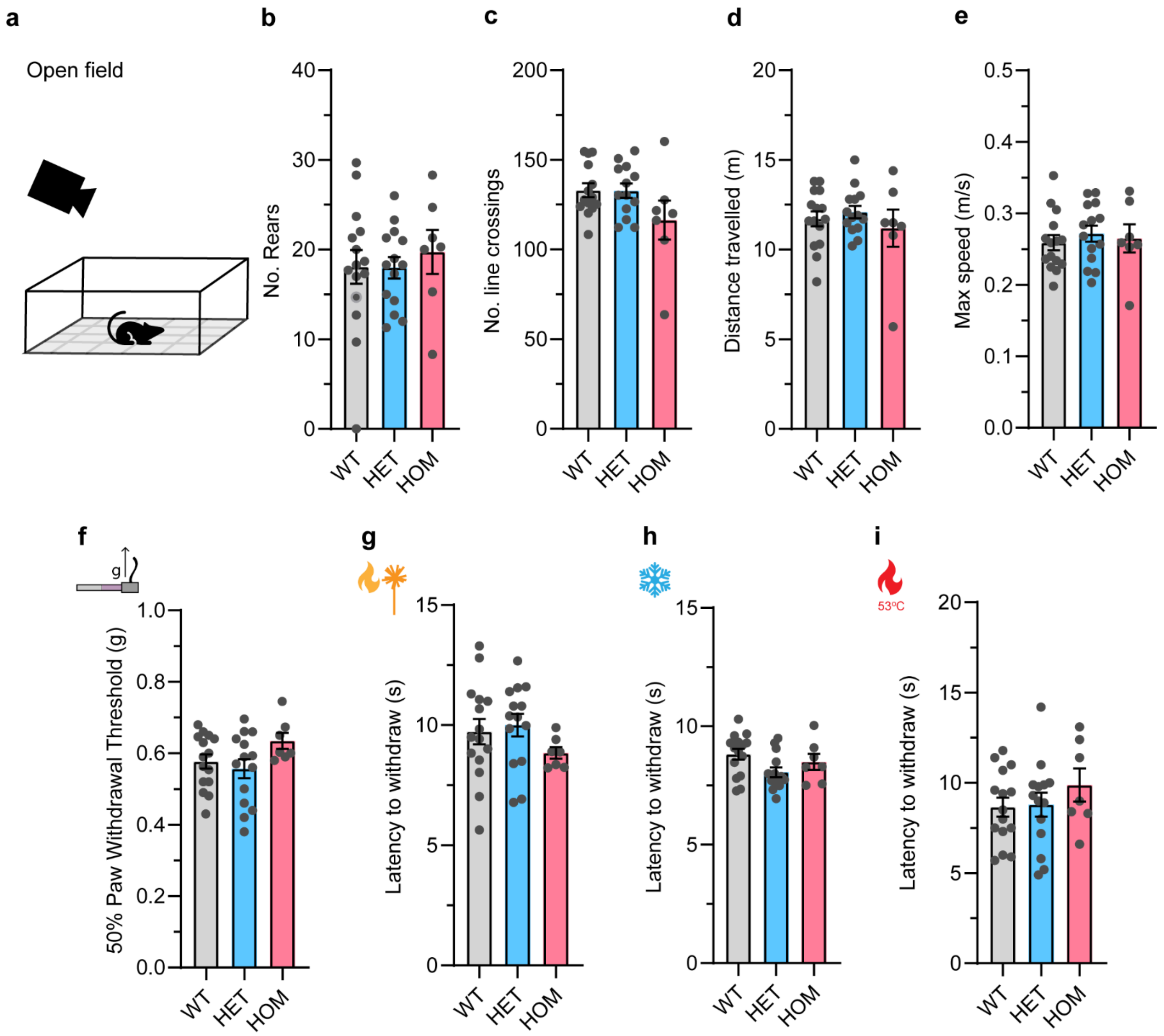
a
Open field
b
No. Rears
0
10
20
30
40
WT
HET
HOM
c
No. line crossings
0
50
100
150
200
WT
HET
HOM
d
Distance travelled (m)
0
5
10
15
20
WT
HET
HOM
e
Max speed (m/s)
0.0
0.1
0.2
0.3
0.4
0.5
WT
HET
HOM
f
g
50% Paw Withdrawal Threshold (g)
0.0
0.2
0.4
0.6
0.8
1.0
WT
HET
HOM
g
Latency to withdraw (s)
0
5
10
15
WT
HET
HOM
h
Latency to withdraw (s)
0
5
10
15
WT
HET
HOM
i
53ºC
Latency to withdraw (s)
0
5
10
15
20
WT
HET
HOM

**Extended data Fig. 8. Behavioural assays that are normal in SLC45A4 HET and KO mice.** **a,** Example of the open field assay where mice are monitored while they explore a chamber with a grid marked into the base. Number of rears **b**, No. of line crossings **c**, distance travelled **d,** and max speed **e**, are all normal in *SLC45A4* HET and KO mice. The withdrawal threshold from a von Frey stimulus **f**, the latency to withdraw from Hargreaves test **g**, the latency to withdraw from a cold stimulus (dry ice) **h,** and the latency to withdraw from a noxious 53$^{o}$C hotplate, were all normal in *SLC45A4* HET and KO mice. WT n = 15 mice, HET n = 14 mice, KO n = 7 mice. one way ANOVA, with Tukey post-hoc test (P > 0.05).

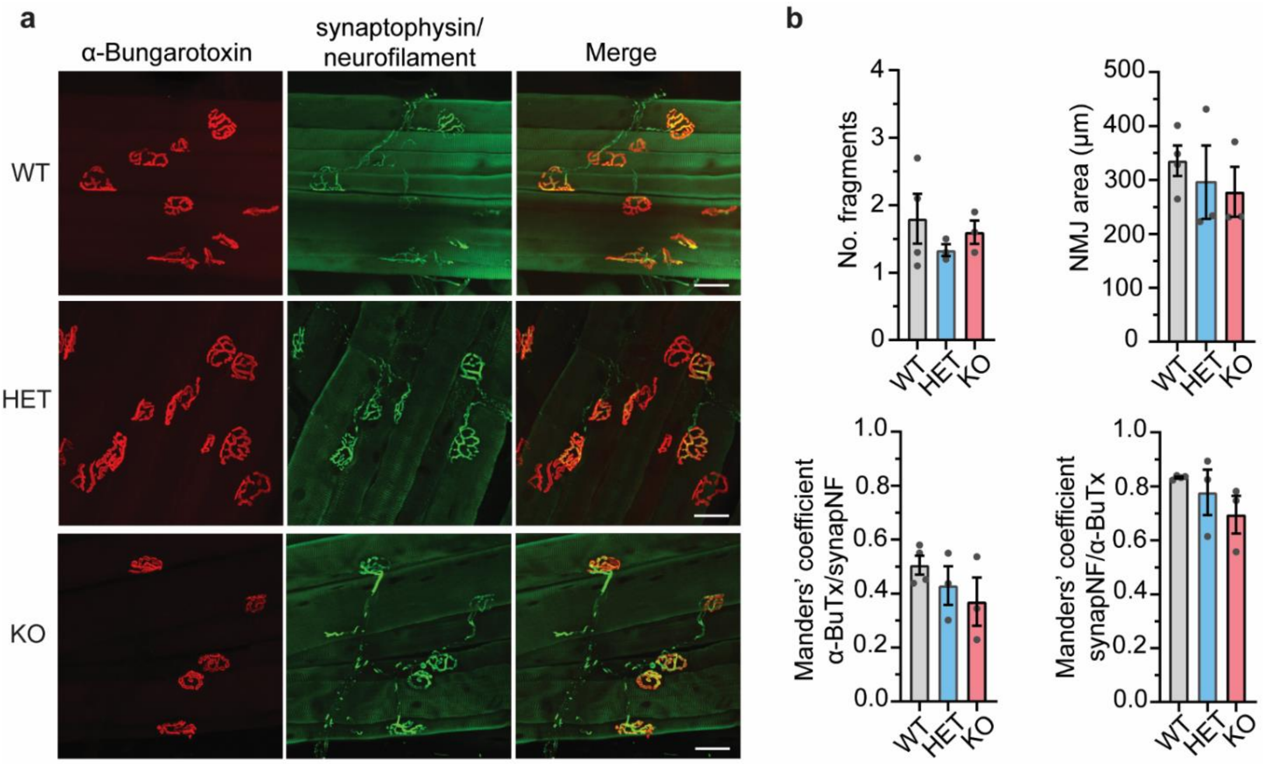

a
α-Bungarotoxin
synaptophysin/ neurofilament
Merge
WT
HET
KO
b
No. fragments
NMJ area (μm)
Manders' coefficient α-BuTx/synapNF
Manders' coefficient synapNF/α-BuTx
WT
HET
KO

**Extended data figure 9: SLC45A4 mutant mice have normal neuromuscular junctions. a,** Example images of neuromuscular junctions (NMJs) from WT, HET and KO mice. Red is α-bungarotoxin, green is synaptophysin & neurofilament. **b,** NJMs were analysed for each genotype, and the no. fragments, NMJ area, co-localisation of α-bungarotoxin/synaptophysin & neurofilament and the co-localisation of synaptophysin & neurofilament/α-bungarotoxin, were all normal. WT n = 4 mice, HET = 3 mice, KO = 3 mice, 10 NMJs were analysed per animal. one way ANOVA, with Tukey post-hoc test, $P > 0.05$. Data mean ± s.e.m, scale bars 40 µm.

#### Extended data tables

| | Data source | rsID | Ch | effect allele（non effect allele） | EAF | Beta | SE | P-value | Position | eQTL | Gene prioritisation (L2G) | CADD |
|---|---|---|---|---|---|---|---|---|---|---|---|---|
| The leading variants | UKB (EU) | rs3905668 | 3 | G (A) | 0.27 | 0.02 | 0.004 | 1.22 x $10^{-8}$ | intergenic | *MSL2, PCCB, PPP2R3A* | *MSL2* (0.28) | 4.25 |
| | | rs10625280 | 8 | TAGAC (T) | 0.59 | 0.018 | 0.003 | 3.37 X $10^{-8}$ | intronic | *SLC45A4, DENND3* | *SLC45A4* (0.49) | 1.79 |
| | | rs3739238 | 8 | C (T) | 0.59 | 0.018 | 0.003 | 3.63 x $10^{-8}$ | exonic | *SLC45A4, DENND3* | *SLC45A4* (0.86) | 13.84 |
| *SLC45A4* variants replications | | | | | | | | | | | | |
| | MVP pain intensity (EU) | rs10625280 | 8 | TAGAC(T) | 0.42 | 0.026 | 0.005 | 1.21 x $10^{-8}$ | | | | |
| | | rs3739238 | 8 | C(T) | 0.41 | 0.026 | 0.005 | 7.24 x $10^{-9}$ | | | | |
| | MVP pain intensity (EU male) | rs10625280 | 8 | TAGAC(T) | 0.42 | 0.027 | 0.005 | 8.18 x $10^{-9}$ | | | | |
| | | rs3739238 | 8 | C(T) | 0.41 | 0.027 | 0.005 | 4.57 x $10^{-9}$ | | | | |
| | FinnGen pain | rs10625280 | 8 | TAGAC(T) | 0.61 | 0.021 | 0.005 | 1.84 x $10^{-5}$ | | | | |
| | FinnGen pain | rs3739238 | 8 | C(T) | 0.61 | 0.021 | 0.005 | 1.99 x $10^{-5}$ | | | | |

**Extended data table 1. Lead variants associated with pain intensity (most bothersome) and replications.** Genome-wide significant top variants from UKB pain intensity (most bothersome) GWAS (European ancestry). The lead variant (rs10625280) and missense variant (rs3739238) of *SLC45A4*. Replications were completed in 1. the MVP pain intensity GWAS (European ancestry), 2. European male pain intensity GWAS, and 3. FinnGen pain GWAS. EAF, effect allele frequencies; SE, standard error; eQTL, expression quantitative trait loci; L2G, locus-to-gene pipeline score; CADD, combined annotation dependent depletion.

| | Small sensory neurons (nociceptors) | | | | | |
|---|---|---|---|---|---|---|
| IB4 binding | IB4-positive | | | IB4-negative | | |
| Genotype (n) | WT (9) | KO (10) | Test | WT (11) | KO (13) | Test |
| Capacitance (pF) | 16.28 ±1.77 | 17.40 ±1.75 | t-test P = 0.65 | 17.27 ±1.60 | 17.27 ±1.41 | t-test P = 0.99 |
| RMP (mV) | -49.05 ±2.42 | -49.74 ±2.25 | MW test P = 0.66 | -53.77 ±2.51 | -52.23 ±1.24 | MW test P = 0.23 |
| Input resistance (mΩ) | 320.7 ±46.53 | 361.1 ±39.64 | MW test P = 0.30 | 394.4 ±33.65 | 368.0 ±45.67 | t-test P = 0.66 |

**Extended data table 2. Passive membrane properties of WT and SLC45A4 KO nociceptors.** Data presented as mean ± s.e.m.

### Supplementary information

#### Supplementary Methods

##### Neuromuscular junction analysis macro

BETTER AREA MACRO:

```
//macro altered by Ulrike Schulze. This macro uses a manually selected image part, subtracts a background,enhances contrast,
// and uses Otsu to make a mask. Output is a summary file and results file with area (in um) and grey value statistics.

roiManager("reset");
run("Add to Manager");
title = getTitle();
path = getDirectory("image");
run("Duplicate...", "title=image-2");
run("8-bit");

//makes rectangle and crops the image
setTool("rectangle");
makeRectangle (524, 148, 266, 308);
waitForUser( "Pause","select rectangle ROI and press ok");
run("Crop");

//subtracts background
run("Duplicate...", "title=orig");
selectWindow("image-2");
run("Subtract Background...", "rolling=60");

//enhances contrast
run("Enhance Contrast...", "saturated=0.001");

//thresholds background subtracted image by using "Intermodes"
run("Duplicate...", "title=area");
```

```
run("Threshold...");
waitForUser( "press ok when ready");
run("Convert to Mask");
run("Despeckle");

// process mask, smooth
selectWindow("area");
run("Open");
run("Median...", "radius=4"); //median smoothes edges a bit
run("Convert to Mask");
rename("area");
waitForUser("use paintbrush to remove extraneous objects");

//measure
run("Set Measurements...", "area mean min perimeter integrated display redirect=[orig] decimal=3");
run("Analyze Particles...", "size=1.25-Infinity circularity=0.00- 1.00 show=Masks display summarize add");
//outlines, bare outlines, ellipses will always include the holes in the structure=> don't use

close("image-2");
close("Mask of area");

dir2 = getDirectory("Output");
selectWindow("Results");
saveAs("Results", dir2+"Results.xls");
selectWindow("Summary");
saveAs("Text", dir2+"Summary.txt");
selectWindow("area");
run("Flatten");
rename("area_image.tif");
saveAs("Tiff", dir2+"area_image.tif");
run("Close");
selectWindow("orig");
saveAs("Tiff", dir2+"orig.tif");
```

#### Supplementary Tables

Supplementary Table 1: Most Bothersome Chronic Pain GWAS Top Results (Excel File)

Supplementary Table 2: PheWAS analyse of secondary phenotypes linked to genetic variants (Excel File)

Supplementary file 1: Validation report: Cryo-EM structure of human SLC45A4 in detergent

Supplementary file 2: Validation report: Cryo-EM structure of human SLC45A4 in lipid nanodiscs

Supplementary tables 3-6 below.

| | *Hs*SLC45A4 LMNG:CHS (EMDB-51377) (PDB 9GIU) | *Hs*SLC45A4 Nanodiscs (EMDB-51365) (PDB 9GHZ) |
|---|---|---|
| **Data collection and processing** | | |
| Magnification | 165,000x | 105,000x |
| Voltage (kV) | 300 | 300 |
| Electron exposure (e–/Å$^2$) | 57.6 | 39.71 |
| Defocus range (μm) | -0.6 to -2.5 | -0.75 to -2.00 |
| Pixel size (Å) | 0.732 | 0.832 |
| Symmetry imposed | C1 | C1 |
| Initial particle images (no.) | 10,040,401 | 20,196,090 |
| Final particle images (no.) | 700,436 | 227,752 |
| Map resolution (Å) | 2.83 | 3.25 |
| FSC threshold | 0.143 | 0.143 |
| Map resolution range (Å) | 2.43-16.59 | 2.91-8.39 |
| **Refinement** | | |
| Initial model used (PDB code) | | 9GIU |
| Model resolution (Å) | 2.83 | 3.25 |
| FSC threshold | 0.143 | 0.143 |
| Model resolution range (Å) | 2.43-16.59 | 2.91-8.39 |
| Map sharpening *B* factor (Å$^2$) | | -163 |
| Model composition | | |
| Non-hydrogen atoms | 4128 | 4307 |
| Protein residues | 493 | 500 |
| Ligands | 8 | 14 |
| Waters | 17 | 12 |
| *B* factors (Å$^2$) | | |
| Protein | 48.06 | 63.25 |
| Ligand | 63.52 | 76.71 |
| Waters | 45.86 | 56.11 |
| R.m.s. deviations | | |
| Bond lengths (Å) | 0.003 | 0.003 |
| Bond angles (°) | 0.534 | 0.538 |
| Validation | | |
| MolProbity score | 1.38 | 1.52 |
| Clashscore | 6.94 | 7.33 |
| Poor rotamers (%) | 0.00 | 0.24 |
| Ramachandran plot | | |
| Favored (%) | 98.36 | 97.37 |
| Allowed (%) | 1.64 | 2.63 |
| Disallowed (%) | 0.00 | 0.00 |

**Supplementary table 3.** CryoEM data collection, refinement and validation statistics.

| | WT | HET | KO | Total |
|---|---|---|---|---|
| No. animals | 31 | 51 | 12 | 94 |
| % animals | 32.9 | 54.3 | 12.8 | 100 |
| No. males | 18 | 24 | 4 | 46 |
| No. females | 13 | 27 | 8 | 48 |
| Male weight (g) | 26.57 ± 0.71 (6) | 26.09 ± 0.88 (7) | 25.43 ± 1.60 (3) | |
| Female weight (g) | 20.70 ± 1.10 (5) | 20.60 ± 1.02 (7) | 20.60 ± 1.47 (5) | |

**Supplementary table 4. Transgenic mouse viability.** Data collected from Het x Het breeding. Fewer KO mice are born than expected. Weights were collected at 8.5-10wks of age. Mean ± SD (n).

| Genotyping Primer | Primer | Sequence | Length |
|---|---|---|---|
| Common Primer | Fwd | TAAAATGGGGAGTCTTGCTGATCTT | 25 |
| Mutant Primer | Rev | TAAAGGCCTGGAAGGTGTGGATT | 23 |
| Wild type Primer | Rev | GACCACTATGTTGCTGGTACTGA | 23 |
| **qPCR Primer** | **Primer** | **Sequence** | **Length** |
| SLC45A4 (exons 4-5) | Fwd | TTCGTGCCTACCTGCTGGATGT | 22 |
| | Rev | GCGTCTGGAACCAGTCACCTA | 21 |
| SLC45A4 (exons 5-6) | Fwd | CTCACTTGGTTCTCCGTCATC | 21 |
| | Rev | ATCTTCACGCCAGCATTGTA | 20 |
| mu HPRT | Fwd | GTCCTGTGGCCATCTGCCTAG | 21 |
| | Rev | TGGGGACGCAGCAACTGACA | 20 |
| mu Beta actin | Fwd | CATTGCTGACAGGATGCAGAAGG | 23 |
| | Rev | TGCTGGAAGGTGGACAGTGAGG | 22 |
| mu GAPDH | Fwd | TGTGTCCGTCGTGGATCTGA | 20 |
| | Rev | TTGCTGTTGAAGTCGCAGGAG | 21 |

**Supplementary table 5. Primers used for genotyping and qPCR**

| Primary Antibody | Source | Identifier |
|---|---|---|
| Rb NeuN (1:500) | Abcam | ab177487 |
| Ms βIII-Tubulin (1:500) | R&D Systems | MAB1195 |
| βIII-Tubulin-FITC conjugated (1:500) | Abcam | ab224978 |
| Sh CGRP (1:500) | Enzo | BML-CA1137 |
| Rb CGRP (1:500) | BMA Biomedicals | T-4032 |
| IB4, streptavidin conjugated (1:100) | Sigma | L2140 |
| Ch NF200 (1:5000) | Abcam | ab4680 |
| Sh TH (1:500) | Millipore | AB1542 |
| Rb TH (1:500) | Millipore | AB152 |
| Ms anti-FLAG | Merck | F1804 |
| Ms anti-β-actin | Merck | A2228 |
| Rb anti-Na+/K+ ATPase | ThermoFisher | ST0533 |
| **Secondary Antibody** | **Source** | **Identifier** |
| Rb PcBl (1:250) | Life Technology, | P-10994 |
| Ms PcBl (1:250) | Thermofisher | P31582 |
| Stp PcBl (1:250) | Life Technology | S11222 |
| Sh Alexa 546 (1:500) | Life Technology | A21098 |
| Rb Alexa 488 (1:500) | Life Technology | A11008 |
| Rb Alexa 546 (1:500) | Life Technology | A11010 |
| Stp Alexa 488 (1:500) | Life Technology | S11223 |
| Ch Alexa 488 (1:500) | Abcam | ab150169 |
| Ch Alexa 546 (1:500) | Life Technology | A11040 |
| Stp Alexa 546 (1:500) | Life Technology | S11225 |
| NeuroTrace (1:10) | Life Technology | N21382 |
| Goat anti-Mouse IgG (H+L) AlexaFluor-488 | ThermoFisher | (A28175) |
| Goat anti-Rabbit IgG (H+L) AlexaFluor-647 | ThermoFisher | (A-21245) |

**Supplementary table 6. Antibodies used in this study**
